# AP2/ERF transcription factor RAP2.6 regulates early flowering in *Arabidopsis thaliana* by altering *S*-nitrosothiol levels and cytokinin responses

**DOI:** 10.64898/2026.05.13.725052

**Authors:** Ashim Kumar Das, Mohammad Golam Mostofa, Da-Sol Lee, Byung-Wook Yun

## Abstract

RAP2.6, an AP2/ERF transcription factor (TF), regulates plant stress responses; however, its role in floral transition remains unexplored. Here, we evaluated RAP2.6’s role in flowering and the associated transcriptional changes in *Arabidopsis thaliana* under long-day conditions. *RAP2.6*-overexpressing line showed early flowering with fewer rosette leaves, whereas *rap2.6-1* mutant flowered later, had more rosette leaves, and higher expression of the floral repressor *FLOWERING LOCUS C* (*FLC*). Early flowering in the overexpressing line was accompanied by transcriptional activation of the floral integrators *GIGANTEA* (*GI*), *FLOWERING LOCUS T* (*FT*), and *COSTANS* (*CO*), potentially through RAP2.6 interaction with GCC/DRE *cis*-regulatory elements. *RAP2.6*-mediated floral transition depended on nitric oxide (NO), with flowering time largely varying based on NO bioactivity. *RAP2.6* was found to be a downstream regulator of *Arabidopsis S-NITROSOGLUTATHIONE REDUCTASE 1* (*GSNOR1*) in controlling S-nitrosothiol (SNO) levels, flowering time, and silique formation. The *NITRIC OXIDE-ASSOCIATED 1* (*NOA1*)-dependent reduction in NO levels abolished early flowering in *35S::RAP2.6* plants without affecting silique formation. Furthermore, enhanced cytokinin sensitivity and upregulation of cytokinin biosynthetic genes suggest cytokinin involvement in RAP2.6-mediated flowering. Together, these findings highlight the crucial role of RAP2.6 in regulating flowering time by integrating redox and hormonal signaling to coordinate reproductive development in *A. thaliana*.

## 1. Introduction

Flowering marks the developmental transition from the vegetative to the reproductive stage and represents one of the tightly regulated processes in plants. In *Arabidopsis thaliana*, flowering time is controlled by multiple endogenous and environmental cues, including vernalization, temperature variation, photoperiod, light quality, plant age, and phytohormonal signaling ^1,2^. Under the photoperiodic pathway, floral transition is orchestrated through coordinated transcriptional regulation of *GIGANTEA (GI)*, *FLOWERING LOCUS T (FT)*, *COSTANS (CO)*, *SUPPRESSOR OF OVEREXPRESSION OF CONSTANS 1 (SOC1)*, and *FRUITFUL (FUL)*, which collectively promote floral induction ^3,4^. Conversely, flowering repressors like *TEMPRANILLO 1 (TEM1)*, *FLOWERING LOCUS C (FLC)*, and *CYCLING DOF FACTOR 1 (CDF1)* antagonize this process by suppressing the expression of early flowering genes ^5,6^. Besides, plant hormones such as gibberellic acid (GA), brassinosteroids (BRs), auxin, jasmonic acid (JA), cytokinin, and abscisic acid (ABA) act as upstream regulators and interact with multiple pathways to coordinate floral transition ^7,8^.

Accumulating evidence indicates that redox signaling also plays a central role in regulating flowering time and its associated downstream transcriptional networks ^9^. Among redox-active signaling molecules, nitric oxide (NO) has emerged as an important regulator of flowering in *A. thaliana* ^10–12^. In plants, NO is primarily generated via the nitrate reduction pathway, in which nitrate reductase (NR) catalyzes the sequential reduction of nitrate to nitrite and subsequently to NO ^13,14^. Although *NITRIC OXIDE-ASSOCIATED 1* (*NOA1*) was initially proposed to function as a nitric oxide synthase (NOS)-like enzyme, subsequent studies identified NOA1 as a circularly permuted GTPase (cGTPase) that indirectly regulates NO homeostasis ^15,16^. A major bioactive reservoir of NO is S-nitrosoglutathione (GSNO), which is formed through the reaction between NO and reduced glutathione (GSH). GSNO serves as a mobile NO donor and mediates protein *S*-nitrosylation by transferring its NO moiety to the sulfhydryl (-SH) group on target proteins, thereby modulating protein activity and signaling ^17,18^. Cellular GSNO and total *S*-nitrosothiol (SNO) levels are tightly controlled by GSNO reductase (GSNOR), which catalyzes GSNO degradation to glutathione disulfide (GSSG) and ammonium (NH ^+^) ^19^.

Genetic evidence supports an important role for GSNOR1-mediated NO homeostasis in flowering regulation. In *Arabidopsis*, the gain-of-function GSNOR1 mutant *gsnor1-1* exhibits delayed flowering under both long- and short-day conditions, whereas the loss-of-function mutant *gsnor1-3*, characterized by elevated GSNO accumulation, flowers earlier than wild-type plants ^11^. These phenotypes are associated with altered expression of key flowering regulators, including *FLC* and *CO*, suggesting that GSNO-dependent redox-signaling regulates plant photoperiod mechanisms. However, these observations contrast with the NO-overproducing mutant *nox1*, which displays late flowering and elevated *FLC* expression via S-nitrosylation of HISTONE DEACETYLASE 5 (HDA5) and HDA6 ^10,12^. This difference suggests that variations in NO bioactivity states may differentially modulate the redox-signaling pathways to control floral transition.

The APETALA2/ethylene-responsive factor (AP2/ERF) superfamily comprises a large group of plant-specific transcription factors (TFs) with essential roles in development, hormone signaling, and stress adaptation ^20,21^. RELATED TO AP2.6 (RAP2.6) belongs to group X of the *Arabidopsis* AP2/ERF family and was initially identified as CORONATINE INSENSITIVE 1 (COI1)-dependent jasmonate-inducible TF ^22,23^. Subsequent studies demonstrated that RAP2.6 participates in responses to ABA, JA, salinity, cold, drought, and pathogen stress ^24–27^. Recently, Zhu et al. ^28^ reported that RAP2.6 physically interacts with CYCLIN-DEPENDENT KINASE 8 (CDK8) and SUCROSE NONFERMENTING 1 (SNF1)-RELATED PROTEIN KINASE 2.6 (SnRK2.6), and directly binds to the GCC/DRE *cis* regulatory elements in stress-responsive genes, namely RESPONSE TO DESICCATION 29A (RD29A) and COLD-REGULATED 15A (COR15A), to mediate drought responses and tolerance. Although several AP2/ERF TFs have been implicated in flowering pattern regulation in rice (*Oryza sativa*), petunia, chrysanthemum (*Chrysanthemum morifolium*), soybean (*Glycine max*), and *Primula forbesii* ^20,29–32^, the developmental function of RAP2.6 in floral transition remains unknown. Notably, NO signaling directly employs AP2/ERF TFs in regulating plant oxygen-sensing mechanisms ^33,34^, raising the possibility that RAP2.6 may function at the intersection of NO signaling and flowering time regulation.

Our earlier RNA-seq analysis identified RAP2.6 among the most strongly upregulated TFs following treatment with 1 mM CysNO, a NO donor ^35,36^, suggesting a potential role in NO-mediated developmental signaling. Based on this observation, we hypothesized that RAP2.6 contributes to NO-dependent regulation of floral transition. Here, we aimed to investigate the role of RAP2.6 in the regulation of flowering time in *A. thaliana* by examining its effects on floral transition and the transcriptional regulation of key flowering genes. We further sought to determine whether RAP2.6-mediated flowering is influenced by cytokinin signaling and GSNOR1-dependent redox homeostasis. Through this work, we aimed to elucidate whether RAP2.6 functions as an integrative regulatory node linking hormonal and redox signaling pathways during reproductive development.

## Results

### GSNO and NO differentially regulate the flowering time in *A. thaliana*

To investigate the impact of differential NO bioactivity on plant floral transition under long-day conditions, transgenic lines with altered GSNO (*gsnor1-3* and *gsnor1-1*) and NO (*cue1-6*, *noa1*, *nia1*, and *nia2*) levels were compared with Col-0 (wild type) (Fig. 1). The number of rosette leaves and bolting time were recorded to assess the floral transition under long-day conditions. *gsnor1-3* plants with elevated GSNO levels exhibited fewer rosette leaf numbers and reduced bolting time, indicating an early-flowering phenotype (Fig. 1A-C), whereas *gsnor1-1* plants with reduced GSNO levels showed comparable effects with Col-0. Contrarily, elevated cellular NO levels in the NO-overproducing mutant *cue1-6* resulted in a late-flowering phenotype, with increased rosette leaves and delayed bolting time (Fig. 1D-F).

**Fig. 1.**
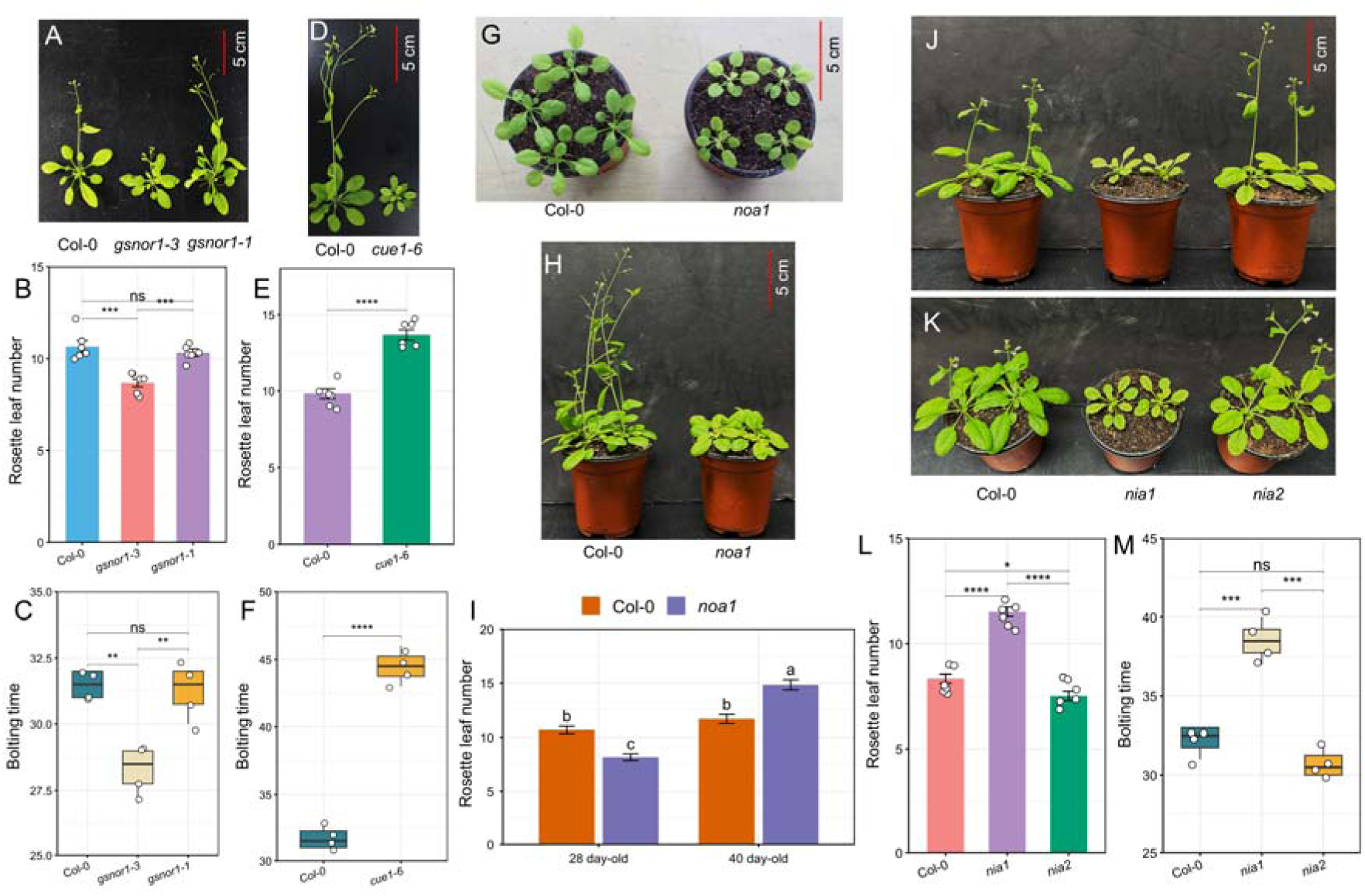
Variation in NO bioactivity differentially regulates flower transition in *Arabidopsis thaliana*. (A-C) Flowering phenotype, rosette leaf number, and bolting time were assessed in 38-day-old soil-grown plants of Col-0, *gsnor1-3*, and *gsnor1-1* under long-day growth conditions. (D-F) Flowering phenotype, rosette leaf number, and bolting time were assessed in 42-day-old soil-grown plants of Col-0 and *cue1-6* under long-day growth conditions. Flowering phenotype and rosette leaf number in (G, I) 28- and (H, I) 40-day-old soil-grown plants of Col-0 and *noa1* under long-day growth conditions. (J-M) Flowering phenotype, rosette leaf number, and bolting time were assessed in 38-day-old soil-grown plants of Col-0, *nia1*, and *nia2* under long-day growth conditions. The data are means ± SE (*n* = 6 for rosette leaf number and *n* = 4 for bolting time). Asterisks indicate significant differences by Student’s two-tailed *t*-test (\**P* < 0.05, \*\**P* < 0.01, \*\*\**P* < 0.001, \*\*\*\**P* < 0.0001). Different letters indicate statistically significant differences by two-way ANOVA and Tukey’s HSD (P < 0.05).

These findings were paralleled with those of Kwon et al. ^11^ and He et al. ^10^. Furthermore, previous reports showed that reduced NO levels in *noa1* led to earlier flower development when compared with Col-0 ^10,12^. Interestingly, we observed fewer rosette leaves in 28-day-old *noa1* plants than in Col-0, consistent with an early-flowering phenotype (Fig. 1G, I). In contrast, this phenotype in soil-grown *noa1* plants disappeared at 40-day-old plants under long-day conditions, showing an increased number of rosette leaves (Fig. 1H, I). To confirm the previous reports, we also sowed *noa1* seeds in MS media and counted the number of rosette leaves at 21 days. However, no notable difference in the number of rosette leaves was observed between Col-0 and *noa1* (Supplementary Fig. 1). We also assessed the flowering time in NR mutants *nia1* and *nia2*, which exhibited distinct patterns of floral transition. Higher rosette leaf number in *nia1* resulted in a late-flowering phenotype, while *nia2* had an early bolting time (Fig. 1J–M). Collectively, these results suggest that variations in NO concentration and regulation exert diverse developmental functions in plants.

### Loss-of-function mutation in RAP2.6 delays flowering time

We first quantified SNO levels in RAP2.6 to correlate with the CysNO-induced upregulation of RAP2.6 observed in our previous RNA-seq analysis ^35,36^. We found that *35S::RAP2.6* plants accumulated higher SNO levels than Col-0, whereas *rap2.6-1* loss-of-function mutant plants had lower SNO levels (Supplementary Fig. 2). Similar to the early-flowering phenotype and increased cellular SNO levels in *gsnor1-3*, we were interested in understanding the functional role of RAP2.6 in the floral transition. We assessed the flowering phenotypes of both the RAP2.6 overexpression and the T-DNA insertion mutant under long-day conditions. Compared to Col-0, the overexpression line of *35S::RAP2.6* exhibited an early-flowering phenotype with decreased rosette leaf number and early bolting (Fig. 2A-C).

**Fig. 2.**
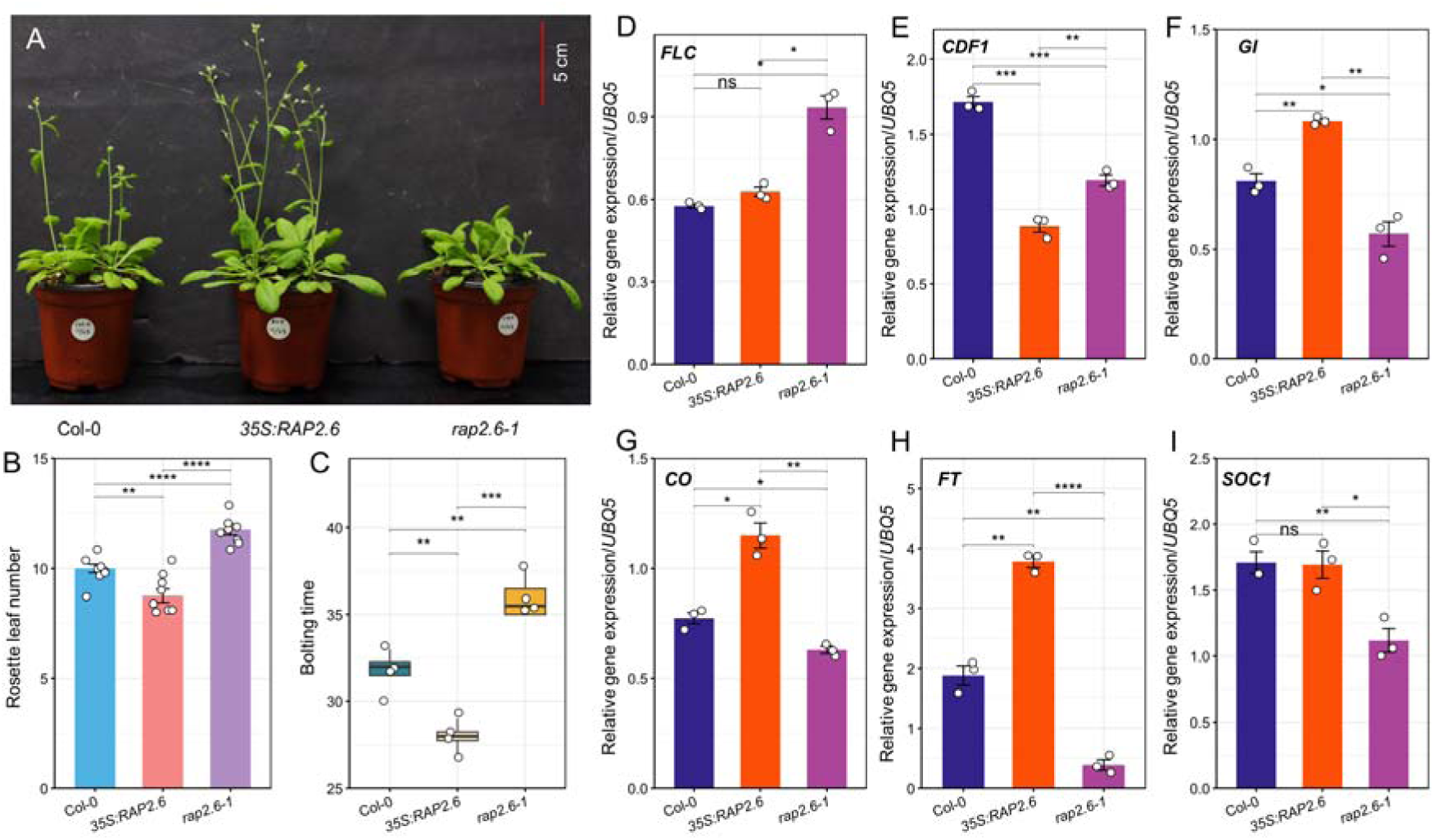
ERF transcription factor RAP2.6 positively regulates the flowering time. (A-C) Flowering phenotype, rosette leaf number, and bolting time were assessed in 38-day-old soil-grown plants of Col-0, *35S::RAP2.6*, and *rap2.6-1* under long-day growth conditions. The data are means ± SE (*n* = 8 for rosette leaf number and *n* = 4 for bolting time). (D-I) Related gene expression analysis of the flowering-time-related marker genes *FLC*, *CDF1*, *GI*, *CO*, *FT*, and *SOC1* in Col-0, *35S::RAP2.6*, and *rap2.6-1* under long-day growth conditions. *UBQ5* was used as an internal control. The data are means ± SE (*n* = 3 biological replicates). Asterisks indicate significant differences by Student’s two-tailed *t*-test (\**P* < 0.05, \*\**P* < 0.01, \*\*\**P* < 0.001, \*\*\*\**P* < 0.0001). FLC, FLOWERING LOCUS C; CDF1, CYCLING DOF FACTOR 1; GI, GIGANTEA; CO, COSTANS; FT, FLOWERING LOCUS T; SOC1, SUPPRESSOR OF OVEREXPRESSION OF CONSTANS 1.

To further confirm the early-flowering phenotypes of *35S::RAP2.6*, we tested the expression of the floral repressor *FLC* and *CDF1*. The transcript level of *FLC* remained comparable to Col-0, while the expression of *CDF1* was markedly reduced in *35S::RAP2.6* plants at bolting (Fig. 2D, E). To understand the functional difference, we also assessed flowering time and associated gene expression in *rap2.6-1* plants. Notably, *rap2.6-1* showed a significant delay in flowering time, as evidenced by a higher number of rosette leaves and later bolting compared to Col-0 (Fig. 2A-C). A significant *FLC* expression further confirmed this late-flowering phenotype in *rap2.6-1* (*P* < 0.5), despite a lower *CDFI* transcript level than in Col-0, indicating that *FLC* controls the flowering time of RAP2.6 (Fig. 2D, E). Furthermore, we evaluated the flower-promoting marker regulators in *35S::RAP2.6* and *rap2.6-1*. Results showed that *35S::RAP2.6*, with an early-flowering phenotype, enhanced transcript levels of *GI*, *CO*, and *FT*, but this upregulation diminished in *rap2.6-1* (Fig. 2F-H).

Transcript level of *SOC1* in *35S::RAP2.6* was comparable to Col-0, while it was significantly reduced in *rap2.6-1* (Fig. 2I). These changes in the relative gene expression levels of flowering-related genes were consistent with the known late-flowering phenotype in *rap2.6-1*, indicating that this developmental defect may result from *FLC*-mediated suppression of *GI*, *CO*, *FT*, and *SOC1*. As RAP2.6 can directly bind to the GCC or DREs of RD29A and COR15A and regulates the transcriptional changes ^24,28^, we explored whether the early flowering gene could be directly regulated by RAP2.6 using *in silico* promoter analyses. *GI*, *CO*, and *FT* promoters contained multiple predicted RAP2.6 binding sites in RD29A and COR15A within 1 kb upstream of the TSS (Supplementary Fig. 3), suggesting that potential transcriptional regulation by RAP2.6 could link to the early flowering phenotype.

### RAP2.6 acts as a downstream regulator of GSNOR1 in floral transition

Based on the increased SNO levels in RAP2.6 overexpressing plants, we hypothesized that early bolting in *35S::RAP2.6* may be linked to GSNOR-dependent flowering mechanisms ^11^. To further explore the GSNOR-mediated role in RAP2.6, we generated new double mutants combining gain- and loss-of-function mutations in RAP2.6 and GSNOR1 to study how GSNO influences flowering through RAP2.6. Results showed that the early-flowering phenotype of *35S::RAP2.6* persisted in the *35S::RAP2.6 gsnor1-3* double mutant, despite the dominant bushiness of elevated GSNO caused by *gsnor1-3* (Supplementary Fig. 4A). Fewer rosette leaves and early bolting were recorded in *35S::RAP2.6* and *35S::RAP2.6 gsnor1-3* compared to Col-0 (Supplementary Fig. 4B, C). Notably, the late-flowering phenotype in *rap2.6-1* had disappeared in both *rap2.6-1 gsnor1-1* and *rap2.6-1 gsnor1-3* plants (Fig. 3A). It is known that *gsnor1-3* shows an early-flowering phenotype compared to both Col-0 and *gsnor1-1*, but no significant differences were observed between Col-0 and *gsnor1-1*, as similar reports by Kwon et al. ^11^ and in the reevaluation of this study. Consequently, plants with fewer rosette leaves and earlier bolting time were observed in *rap2.6-1 gsnor1-3*, but *gsnor1-3*-mediated overdominance in bushiness and silique size (Fig. 3A-C; Supplementary Fig. 5). However, the strong late-flowering phenotype of *rap2.6- 1* was also rescued by *gsnor1-1* in *rap2.6- 1 gsnor1- 1*, as shown by similar rosette leaf numbers and bolting times as in Col-0 (Fig. 3A-C).

**Fig. 3.**
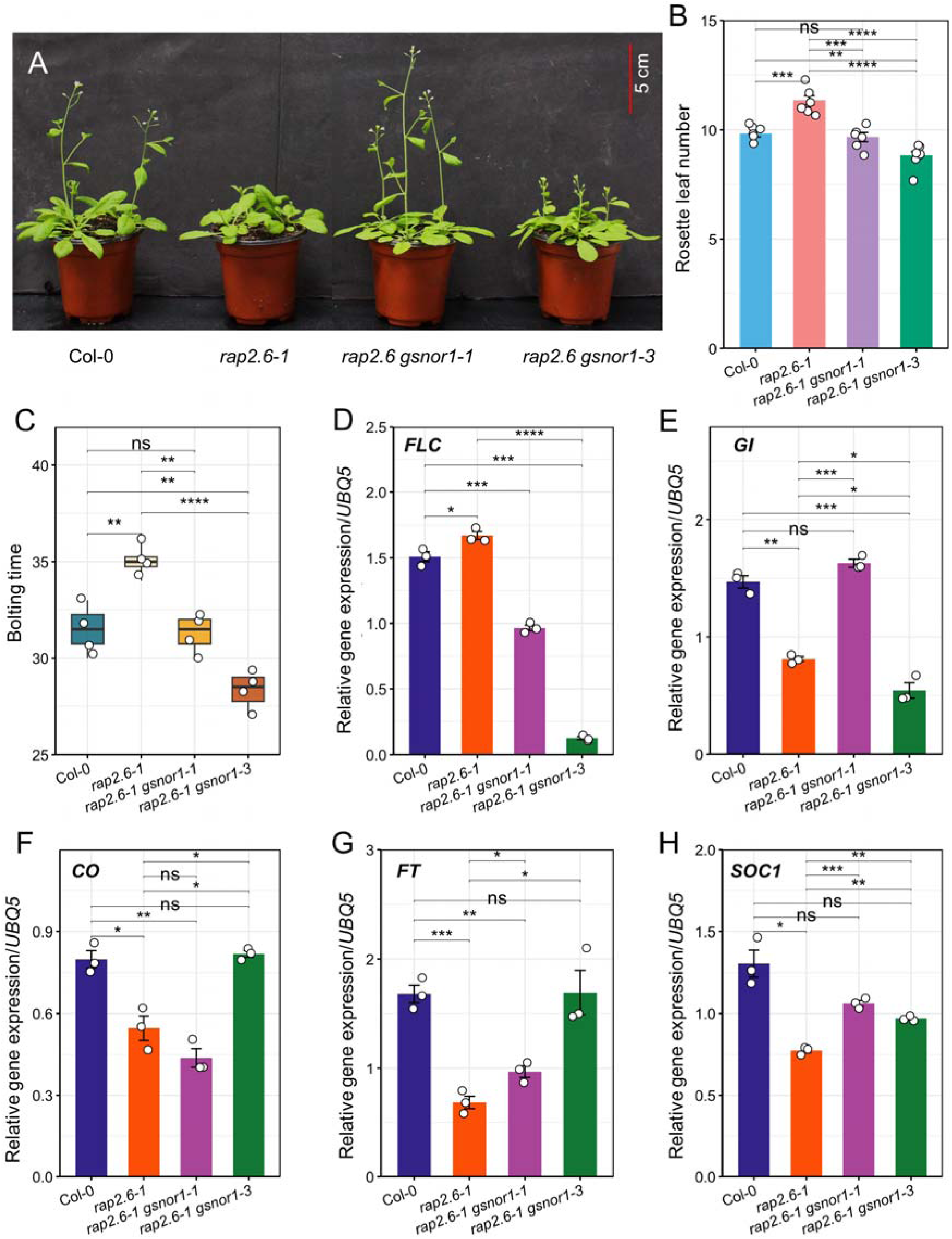
Altering GSNO levels affects flowering time in the RAP2.6 loss-of-function mutation. (A-C) Flowering phenotype, rosette leaf number, and bolting time were assessed in 38-day-old soil-grown plants of Col-0, *rap2.6-1*, *rap2.6-1 gsnor1-1*, and *rap2.6-1 gsnor1-3* under long-day growth conditions. The data are means ± SE (*n* = 6 for rosette leaf number and *n* = 4 for bolting time). (D-H) Related gene expression analysis of the flowering-time-related marker genes *FLC*, *GI*, *CO*, *FT*, and *SOC1* in Col-0, *rap2.6-1*, *rap2.6-1 gsnor1-1*, and *rap2.6-1 gsnor1-3* under long-day growth conditions. *UBQ5* was used as an internal control. The data are means ± SE (*n* = 3 biological replicates). Asterisks indicate significant differences by Student’s two-tailed *t*-test (\**P* < 0.05, \*\**P* < 0.01, \*\*\**P* < 0.001, \*\*\*\**P* < 0.0001). FLC, FLOWERING LOCUS C; GI, GIGANTEA; CO, COSTANS; FT, FLOWERING LOCUS T; SOC1, SUPPRESSOR OF OVEREXPRESSION OF CONSTANS 1.

To further confirm the GSNOR1-driven rescue of flowering in *rap2.6-1*, we measured the relative expression of flowering-related genes. The relative expression of floral repressor *FLC* was significantly downregulated in *rap2.6-1 gsnor1-3*, consistent with the early-flowering phenotype and the nature of *gsnor1-3* (*P* < 0.01) (Fig. 3D). Similar to Fig. 2D, *FLC* gene expression was notably higher in *rap2.6-1* than in Col-0, whereas this expression remained reduced in *rap2.6-1 gsnor1-1*. Moreover, we compared the expression levels of floral enhancer genes (*GI*, *CO*, *FT*, and *SOC1*) in *rap2.6-1* and the double mutants *rap2.6-1 gsnor1-1* and *rap2.6-1 gsnor1-3* (Fig. 3E-H). The relative expression of *CO* and *FT* significantly increased in *rap2.6-1 gsnor1-3* compared to *rap2.6-1* alone, but remained comparable with Col-0 (*P* < 0.0001). We observed that despite the floral enhancer genes not being upregulated in *rap2.6-1*, the relative expression of *GI* and *SOC1* between *rap2.6-1* and *rap2.6-1 gsnor1-3* showed similar effects, which was unexpected. Moreover, the expression of *GI* and *SOC1* increased in *rap2.6-1 gsnor1-1*, while the expression of *CO* and *FT* decreased, compared to *rap2.6-1 gsnor1-3* (Fig. 3F-G). Overall, our results suggest that the floral rescue of *rap2.6-1* by GSNOR1 is presumably mediated by *FLC*, *CO*, and *FT*, rather than *GI* and *SOC1*.

### Reduced NO levels negatively regulate flowering time in RAP2.6

To further understand how reduced NO availability influences floral transition in the RAP2.6 regulatory pathway, we investigated the genetic interaction between *RAP2.6* and *NOA1*, given that the low-NO mutant *noa1* exhibited delayed flowering (Fig. 1H, I). We generated the double mutant *35S::RAP2.6 noa1* and *rap2.6-1 noa1*. Under long-day conditions, both double mutants showed phenotypes consistent with the individual *noa1* mutant. Intriguingly, the late-flowering phenotype of *noa1* persisted in *35S::RAP2.6 noa1* plants, as demonstrated by a higher number of rosette leaves and late bolting (Fig. 4A-C). Furthermore, the late-flowering phenotype in *noa1* and *35S::RAP2.6 noa1* plants was confirmed by increased transcript levels of the floral repressor *FLC* (Fig. 4D). Although *35S::RAP2.6* showed elevated expression of *CO*, *FT*, and *SOC1*, and were consistent with its early-bolting traits, these responses were significantly reduced in *35S::RAP2.6 noa1* plants (Fig. 4E-G).

**Fig. 4.**
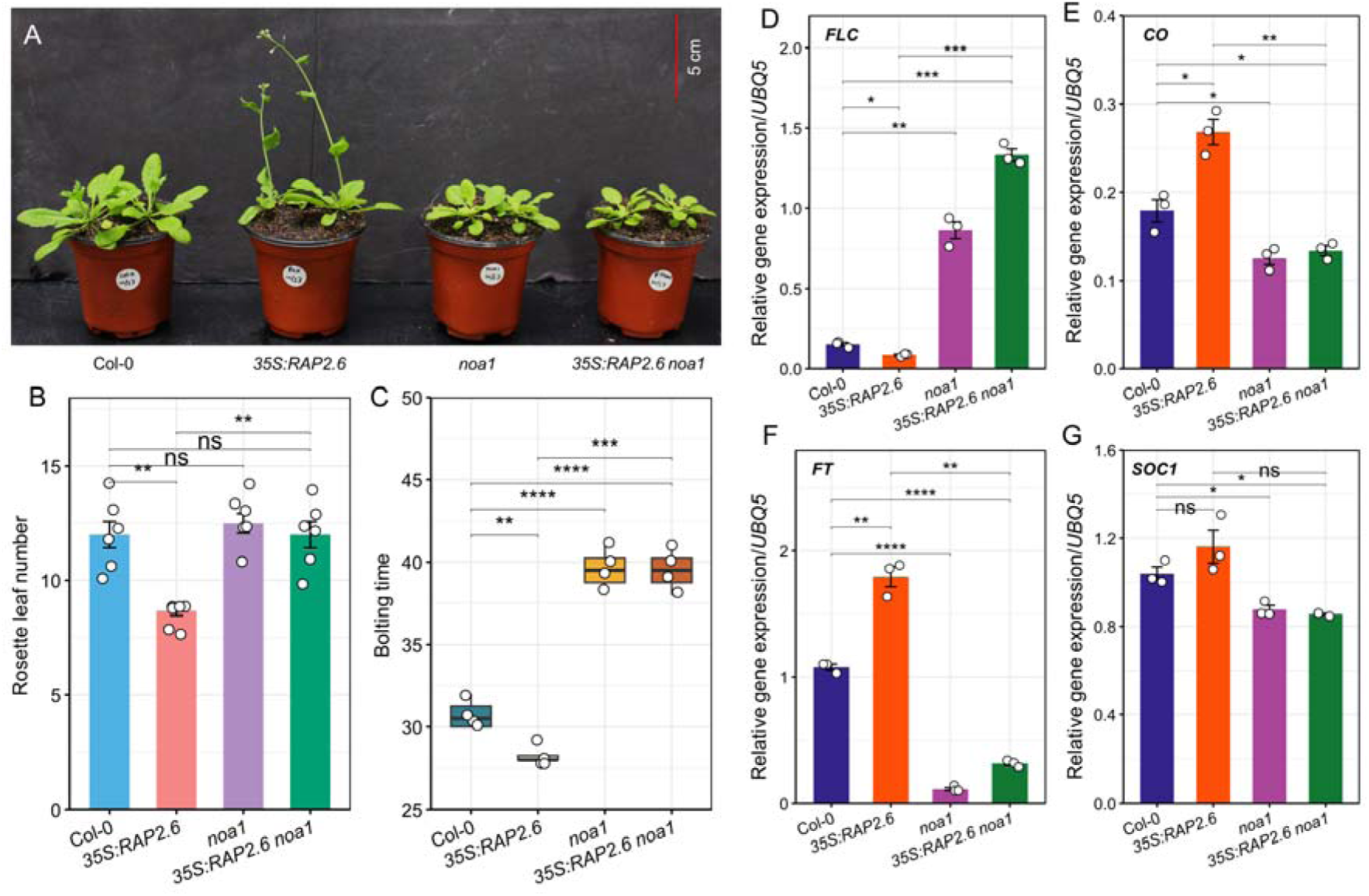
NOA1-dependent reduced NO levels delay flowering time in *35S::RAP2.6*. (A-C) Flowering phenotype, rosette leaf number, and bolting time were assessed in 38-day-old soil-grown plants of Col-0, *35S::RAP2.6*, *noa1*, and *35S::RAP2.6 noa1* under long-day growth conditions. The data are means ± SE (*n* = 6 for rosette leaf number and *n* = 4 for bolting time). (D-H) Related gene expression analysis of the flowering-time-related marker genes *FLC*, *CO*, *FT*, and *SOC1* in Col-0, *35S::RAP2.6*, *noa1*, and *35S::RAP2.6 noa1* under long-day growth conditions. *UBQ5* was used as an internal control. The data are means ± SE (*n* = 3 biological replicates). Asterisks indicate significant differences by Student’s two-tailed *t*-test (\**P* < 0.05, \*\**P* < 0.01, \*\*\**P* < 0.001, \*\*\*\**P* < 0.0001). FLC, FLOWERING LOCUS C; CO, COSTANS; FT, FLOWERING LOCUS T; SOC1, SUPPRESSOR OF OVEREXPRESSION OF CONSTANS 1.

The defect in flowering time conferred by *noa1* remained unchanged in *rap2.6-1* (Fig. 5A-C). The *rap2.6-1 noa1* double mutant also showed increased *FLC* expression accompanied by reduced transcript levels of *CO, FT,* and *SOC1* (Fig. 5D-G). Overdominance of *noa1* is evident not only in flowering but also in their overall phenotypes, as both *35S::RAP2.6 noa1* and *rap2.6-1 noa1* plants closely resembled *noa1*. In contrast, *noa1* had no detectable effect on silique development in either genetic background (Supplementary Fig. 6A, B).

**Fig. 5.**
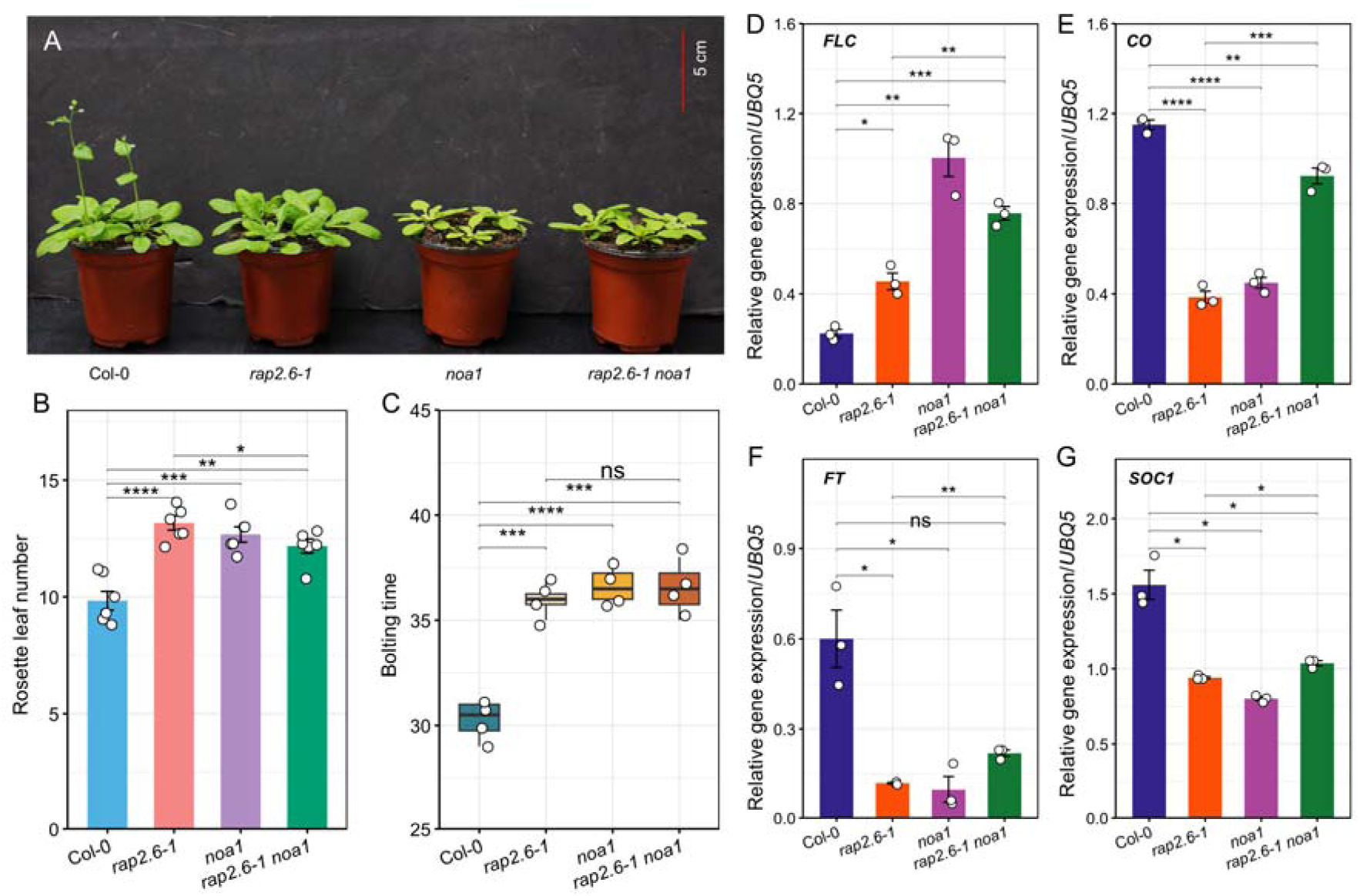
The parallel effect of flowering time between *noa1* and a loss-of-function mutation in RAP2.6. (A-C) Flowering phenotype, rosette leaf number, and bolting time were assessed in 38-day-old soil-grown plants of Col-0, *rap2.6-1*, *noa1*, and *rap2.6-1 noa1* under long-day growth conditions. The data are means ± SE *n* = 6 for rosette leaf number and *n* = 4 for bolting time). (D-H) Related gene expression analysis of the flowering-time-related marker genes *FLC*, *CO*, *FT*, and *SOC1* in Col-0, *rap2.6-1*, *noa1*, and *rap2.6-1 noa1* under long-day growth conditions. *UBQ5* was used as an internal control. The data are means ± SE (*n* = 3 biological replicates). Asterisks indicate significant differences by Student’s two-tailed *t*-test (\**P* < 0.05, \*\**P* < 0.01, \*\*\**P* < 0.001, \*\*\*\**P* < 0.0001). FLC, FLOWERING LOCUS C; CO, COSTANS; FT, FLOWERING LOCUS T; SOC1, SUPPRESSOR OF OVEREXPRESSION OF CONSTANS 1.

To further validate the *noa1*-dependent flowering response, seedlings of *noa1*, *35S::RAP2.6 noa1*, and *rap2.6-1 noa1* were grown on MS medium alongside Col-0, as shown in Supplementary Fig. 1. Under these conditions, we observed no consistent differences in the number of rosette leaves in *35S::RAP2.6 noa1* and *rap2.6-1 noa1* plants (Supplementary Fig. 7). Collectively, these findings demonstrate that reduced NO accumulation in *noa1* exerts a dominant effect on floral transition, overriding RAP2.6-mediated flowering regulation and highlighting the essential role of NO signaling in controlling reproductive development.

### Cytokinin controls early flowering in RAP2.6

Previous studies have shown that *RAP2.6* responds to multiple phytohormones that are known to influence plant growth, development, and flowering ^24,25^. This prompted us to investigate whether the flowering phenotype associated with RAP2.6 is hormonally regulated. We used the *Arabidopsis* eFP Browser to examine the expression response of RAP2.6 to different plant growth hormones, including auxin (IAA), GA (GA-3), cytokinin (zeatin), and BR, all of which are directly involved in the floral transition ^8^ (Supplementary Fig. 8).

Fascinatingly, *RAP2.6* exhibited a stronger transcriptional response to cytokinin during the rosette stage compared with other hormone treatments (Fig. 6A). Given the established role of cytokinin in promoting floral transition and regulating flowering-associated pathways ^37,38^, we further investigated the involvement of cytokinin in RAP2.6-mediated flowering regulation. We evaluated the transcript levels of *CYP735A1* and *CYP735A2*, which encode cytochrome P450 monooxygenases (P450s) responsible for the biosynthesis of *t*Z-type cytokinin ^39^. The qRT-PCR data revealed that both *CYP735A1* and *CYP735A2* were markedly upregulated in *35S::RAP2.6*, whereas downregulated in *rap2.6-1* (Fig. 6B, C). We further examined the expression of *CYTOKININ RESPONSE FACTOR 2* (*CRF2*), an AP2/ERF TF involved in cytokinin signaling ^40^. Although *CRF2* expression was downregulated in *rap2.6-1*, the relative expression in *35S::RAP2.6* was similar to Col-0 (Fig. 6D). To determine whether RAP2.6 influences cytokinin sensitivity, we performed cytokinin response assays and observed that *35S::RAP2.6* seedlings displayed enhanced sensitivity to cytokinin treatment relative to Col-0 plants (Fig. 6E–G). Together, these findings suggest that RAP2.6-mediated floral transition is closely associated with cytokinin biosynthesis and signaling, supporting a cytokinin-dependent mechanism underlying the early-flowering phenotype.

**Fig. 6.**
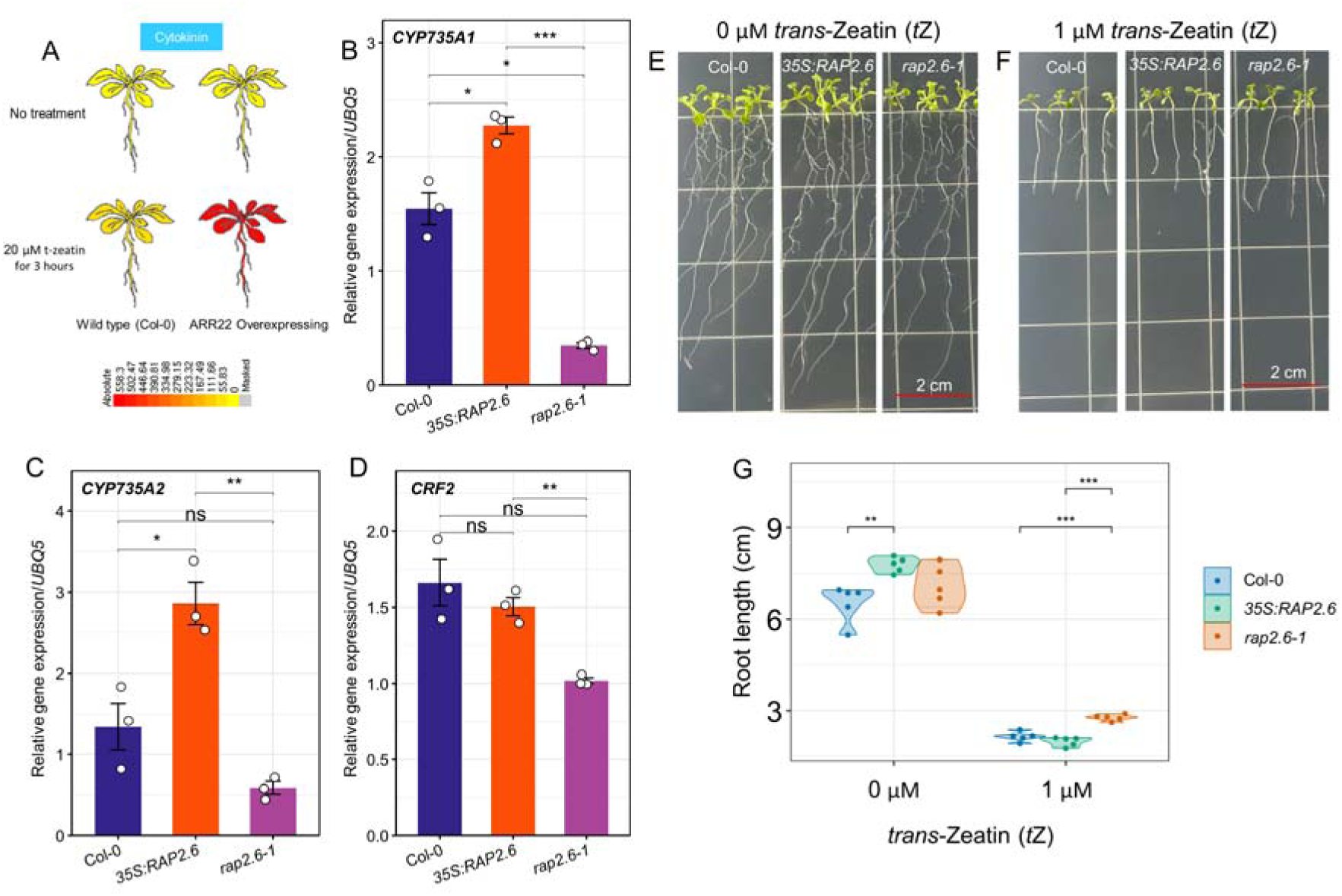
Cytokinin-dependent early flowering in RAP2.6. (A) Visualization of the expression levels of *AtRAP2.6* in the pretreatment with 20 µM t-zeatin, retrieved from *Arabidopsis* eFP Browser. (B-D) Related gene expression analysis of the cytokinin-responsive genes *CYP735A1*, *CYP735A2*, and C*RF2* in Col-0, *35S::RAP2.6*, and *rap2.6-1* under long-day growth conditions. *UBQ5* was used as an internal control. The data are means ± SE (*n* = 3 biological replicates). (E-G) Primary root length of Col-0, *35S::RAP2.6*, and *rap2.6-1* seedlings grown on 1/2 MS media supplemented with 0 and 1 μM *t*Z-zeatin at the post-germination stage. The data are means ± SE (*n* = 8 biological replicates). Asterisks indicate significant differences by Student’s two-tailed *t*-test (\**P* < 0.05, \*\**P* < 0.01, \*\*\**P* < 0.001, \*\*\*\**P* < 0.0001). CRF2, CYTOKININ RESPONSE FACTOR 2.

**Fig. 7.**
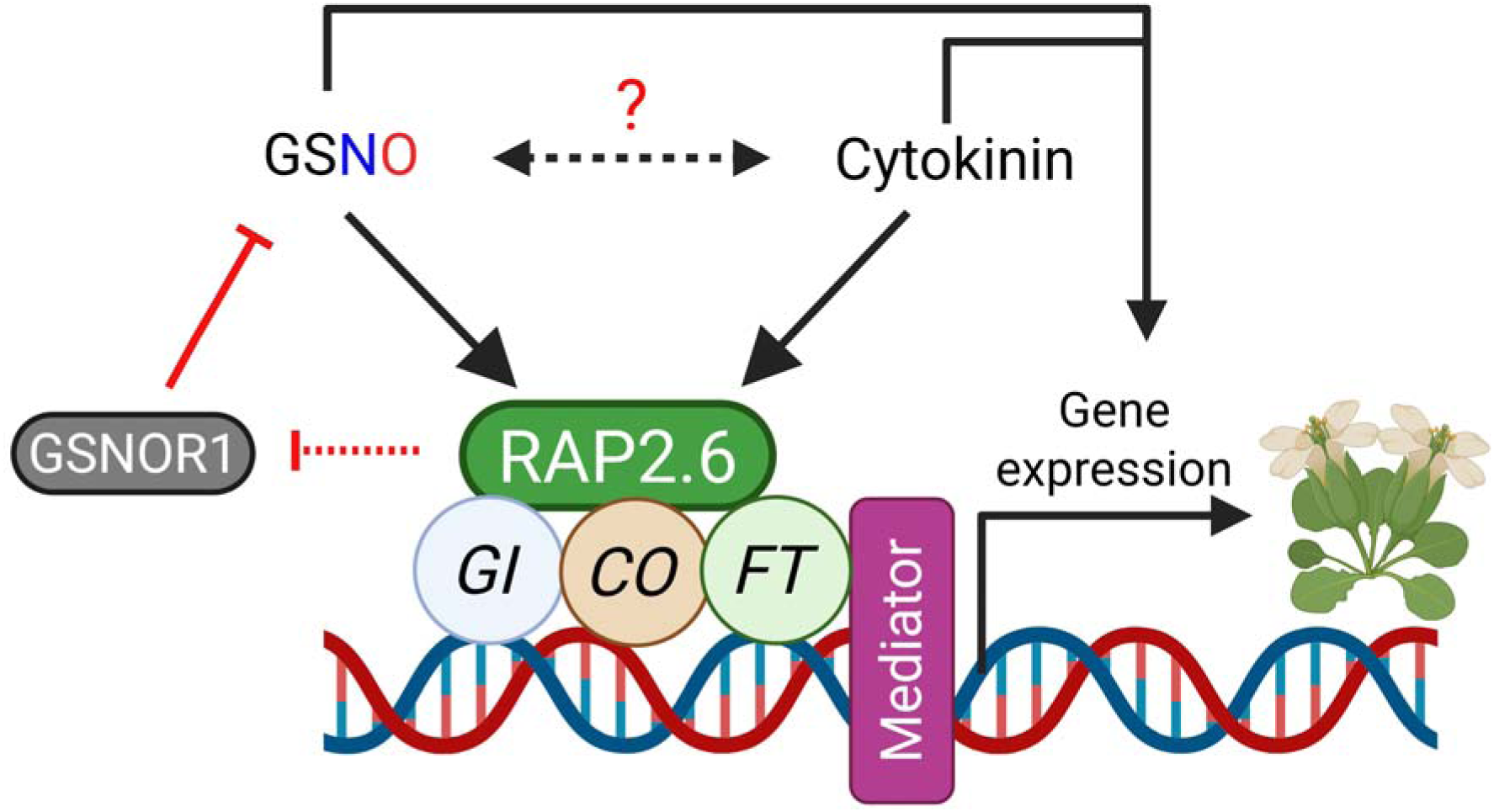
A proposed working model of RAP2.6-mediated early flowering phenotype. Elevated SNO levels in RAP2.6 show a parallel early flowering time to that of *gsnor1-3* (elevated GSNO/SNO level), which also rescues the floral transition of the loss-of-function mutation in *RAP2.6*. RAP2.6 can bind to the promoters of floral activating genes containing GCC/DRE-motif, such as RD29A and COR15A, and promote their expression. Moreover, RAP2.6 promotes flowering time in a cytokinin-dependent manner. However, the mechanism by which GSNO and cytokinin intersect during the floral transition remains to be investigated.

## Discussion

Nitric oxide (NO) is a versatile signaling molecule that regulates various developmental and physiological processes in plants, including developmental transition from vegetative to reproductive stage. The role of NO in flowering was first demonstrated in *A. thaliana*, where the NO-overproducing mutant *nox1*, later identified as an allele of *CUE1*, exhibited a delayed flowering phenotype ^10^. A similar effect was observed in *cue1-6*, a strong allele generated by ethylmethanesulfonate (EMS) mutagenesis, where genetic epistasis showed that the *flc-3* mutant rescued the late-flowering phenotype of *cue1-6* ^41^. Mechanistically, elevated NO promotes *S*-nitrosylation of HDA5 and HDA6, thereby activating *FLC* expression and delaying floral transition ^12^. Because *S*-nitrosylation dynamically modulates cellular redox homeostasis and protein activity ^17,18^, perturbation of SNO metabolism is expected to have profound developmental consequences.

Consistent with this notion, loss of *GSNOR1*, a central regulator of protein de-nitrosylation and GSNO turnover, leads to elevated SNO accumulation and altered plant growth ^42^. Furthermore, Kwon et al. ^11^ reported that the *gsnor1-3* loss-of-function mutant showed an early-flowering phenotype, accompanied by reduced *FLC* and enhanced *CO* transcript levels. In agreement with these findings, our study confirmed distinct flowering phenotypes associated with altered NO bioactivity (Fig. 1). Notably, we observed that the reduced-NO mutant *noa1* displayed delayed flowering under long-day conditions, characterized by increased rosette leaf number and delayed bolting (Fig. 1). Although earlier studies described an accelerated floral transition in *noa1* during early developmental stages ^10^, our observations at late-growth phase indicates that sustained NO deficiency in *noa1* ultimately delays reproductive transition (Fig. 1). These findings suggest that developmental timing and NO bioactivity may differentially influence flowering outputs.

Seligman et al. ^43^ reported an early-flowering phenotype in the *nia1 nia2* double mutant, while the individual roles of *nia1* and *nia2* in flowering had not been reported previously. In this study, we observed a differential floral transition among NR mutants. While *nia1* showed a late-flowering phenotype, *nia2* flowered relatively sooner than Col-0 (Fig. 1). Given that NIA2 accounts for nearly 90% of plant NR activity and is predominantly active in meristematic tissues ^44,45^, its dominant enzymatic contribution may compensate for NIA1 deficiency during floral transition. Differential regulation of *nia1* and *nia2* was also reported in the context of ABA sensitivity and heat tolerance ^46,47^. Collectively, these results support the idea that quantitative and spatial differences in NO production differentially regulate flowering time in plants.

Genetic studies have uncovered multiple regulatory mechanisms underlying the flowering time in *A. thaliana*. Several AP2/ERF members have been directly implicated in the control of flowering. In *Arabidopsis*, RNAi-mediated suppression of *ERF110* significantly delayed bolting ^48^, whereas heterologous expression of *CmERF110* from chrysanthemum promoted early flowering, reduced rosette leaf number, and enhanced expression of key floral regulators, including *CO, FT, SOC1, LFY,* and *FUL*. In contrast, overexpression of *ERF1* delayed bolting by suppressing *FT* expression ^49^, highlighting the functional diversity of ERF proteins in flowering regulation. Similar regulatory roles have been reported in rice, where AP2/ERF TFs such as INDETERMINATE SPIKELET 1 (IDS1), SUPERNUMERARY BRACT (SNB), LATE FLOWERING SEMI-DWARF (LFS), and LEAFY COTYLEDON2 and FUSCA3-LIKE 1 (LFL1) modulate floral organ development and flowering time, thereby affecting yield-related traits (He et al., 2025). Likewise, in soybean, the AP2/ERF TF TIME OF FLOWERING 13B (TOE4b), identified through the genome-wide association study, negatively regulates flowering, as RNAi lines flowered earlier, whereas overexpression delayed flowering and improved yield performance ^32^. These findings provide significant insights into the role of AP2/ERF TFs in plant development. Despite the identification of 122 ERF family members in *Arabidopsis* ^22^, only a limited number have been functionally characterized in flowering regulation. Given the strong NO responsiveness of RAP2.6 and its association with elevated SNO accumulation, we sought to determine whether this stress-responsive AP2/ERF TF also contributes to the regulation of floral transition.

Under the photoperiodic pathway, the floral transition coordinates transcriptional regulation of the florigen protein FT, which is localized in the leaf ^50,51^. CO is a zinc-finger transcription factor, primarily known for flowering activation, binds directly to the FT promoter, and enhances its expression ^52^. FT also moves smoothly and can localize to the shoot apical meristem (SAM), where it forms a complex with the bZIP TF FD. These complex triggers floral transition by activating SOC1 and FUL genes ^53,54^. Additionally, GI controls circadian rhythms and flowering timing, and plants that overexpress GI (*35S:GI*) show increased levels of CO and FT ^55^. We demonstrated that RAP2.6, a group X ERF TF, responds to NO, ABA, JA, salt, cold, drought, and biotic stress ^24–27,36^, and also showed an early-flowering phenotype in *Arabidopsis 35S::RAP2.6* (Fig. 2). These phenotypes were accompanied by reciprocal transcriptional changes in key flowering regulators, with increased expression of *GI, CO, FT,* and *SOC1* in overexpression lines and elevated *FLC* expression in *rap2.6-1* (Fig. 2). These findings strongly support a role for RAP2.6 in promoting floral transition through activation of the photoperiodic flowering pathway. *In silico* analysis also predicted that RAP2.6 plausibly binds the promoter regions of floral enhancer genes (Supplementary Fig. 3), suggesting transcriptional activation.

Earlier studies reported that group VII ERF TFs are largely regulated by NO ^33,34^. We also aimed to examine how redox modification in RAP2.6 alters floral transitions. We found distinct variation in cellular SNO levels between *35S::RAP2.6* and *rap2.6-1* plants (Supplementary Fig. 2), indicating a plausible interaction with NO signaling that controls flowering in RAP2.6. Our newly generated double mutant *35S::RAP2.6 gsnor1-3* retained an early-flowering phenotype consistent with the independent effects of both parental lines (Supplementary Fig. 4). When *gsnor1-3* was considered to flower faster than Col-0, the gain-of-function mutant *gsnor1-1* showed a comparable phenotype to Col-0. More importantly, both *gsnor1-1* and *gsnor1-3* substantially rescued the delayed flowering phenotype of *rap2.6-1*, accompanied by reduced *FLC* expression (Fig. 3B, D). Flowering enhancer genes *CO* and *FT* in *rap2.6-1 gsnor1-3* were largely rescued, indicating visible floral regulation that accelerated flowering time. *gsnor1-1* influenced flowering time in *rap2.6-1 gsnor1-1* plants and showed upregulation of *GI* expression comparable with Col-0, while *CO*, *FT*, and *SOC1* were downregulated, suggesting *gsnor1-1* may not largely follow this pathway. Kwon et al. ^11^ also reported slightly lowered *CO* expression in *gsnor1-1* compared to Col-0 but comparable *FLC* and LFY expression. However, the abnormal silique development observed in *gsnor1-3*-containing double mutants (Supplementary Fig. 5) suggests that excessive SNO accumulation may uncouple flowering acceleration from normal reproductive development.

We also examined how the NOA1-dependent reduction in NO levels affects flowering time in RAP2.6. With the evidence of late-flowering phenotype in *noa1* plants (Fig. 1H, I), our newly generated double mutants of *35S::RAP2.6 noa1* and *rap2.6-1 noa1* plants displayed a parallel phenotype of *noa1* (Fig. 4 and 5). An increased number of rosette leaves and late bolting time were observed in both double mutant plants, which was consistent with the upregulation of flowering repressor *FLC* and downregulation of flowering enhancer *CO*, *FT*, and *SOC1*. Although the contribution of NR-derived NO to RAP2.6 function remains to be clarified, these findings establish NO as an essential upstream determinant of RAP2.6-mediated flowering regulation.

Phytohormones are major regulators of flowering, and RAP2.6 has previously been linked to multiple hormone-responsive pathways. Our data revealed that *RAP2.6* was particularly responsive to cytokinin during the rosette stage (Fig. 6), prompting us to investigate cytokinin metabolism during floral transition. We found that major cytokinin biosynthetic genes *CYP735A1* and *CYP735A2* were notably upregulated in *35S::RAP2.6* plants, and downregulated in *rap2.6-1* (Fig. 6). Moreover, CRF2, an AP2/ERF TF involved in cytokinin signaling, was significantly reduced in *rap2.6-1* (*P* < 0.01) (Fig. 6). Physiological assays further showed enhanced sensitivity of *35S::RAP2.6* seedlings to *tZ* treatment (Fig. 6).

Because elevated cytokinin levels have recently been shown to accelerate flowering by reducing rosette leaf number ^38^, our results suggest that RAP2.6 promotes floral transition, at least in part, through modulation of cytokinin biosynthesis and signaling.

In summary, our study identifies RAP2.6 as a previously unrecognized positive regulator of flowering in *A. thaliana*. *RAP2.6* overexpression exhibited an early-flowering phenotype under long-day conditions through the upregulation of key floral activators, including *GI*, *CO*, and *FT*, whereas loss of *RAP2.6* delayed floral transition and increased *FLC* expression.

Genetic analyses further demonstrated that RAP2.6-mediated flowering is strongly influenced by GSNOR1-dependent SNO accumulation and NO availability, suggesting NO signaling as an essential upstream regulator of RAP2.6 function. In addition, enhanced cytokinin responsiveness and transcriptional activation of cytokinin biosynthetic genes indicate that cytokinin contributes to RAP2.6-dependent floral induction. Together, these findings establish RAP2.6 as an integrative regulatory node linking redox and hormonal signaling during reproductive development. Given its previously characterized roles in abiotic and biotic stress responses, RAP2.6 may represent a promising molecular target for engineering developmental plasticity and yield resilience in crop species.

## Materials and methods

### Plant materials and growth conditions

All allelic mutants and transgenic lines used in this study were on a Columbia (Col-0) ecotype background. Seeds of the T-DNA insertion line of *rap2.6-1* (N301757) and the overexpression line *35S::RAP2.6* were kindly provided by Dr. Holger Bohlmann (University of Natural Resources and Life Sciences, Vienna, Austria) and have been described previously ^27^. The T-DNA mutants *gsnor1-1* and *gsnor1-3*, carrying gain- and loss-of-function mutations in *GSNOR1*, respectively, were reported previously ^56^. The seeds of *cue1-6* (CS3168) and *noa1* (CS6511) used in this study were described previously ^10,15^, and *nia1* and *nia2* were obtained as previously reported ^57^. Double homozygous mutants were obtained by crossing *35S::RAP2.6* or *rap2.6-1* with *gsno1-1*/*gsnor1-3*/*noa1*. Following confirmation of heterozygous F1 plants, self-crossed F2 progeny were genotyped by PCR to identify double homozygous lines, and F3/F4 generation seeds were used for subsequent analyses (Supplementary Fig. 9; Supplementary Table 1). Seeds were allowed to germinate in horticultural soil (Bio Top Soil No.1, Heungnong Trading Company, South Korea) and grow at 22 ± 1°C under long-day conditions (16 h light/8 h dark). Two-week-old seedlings were subsequently transplanted into individual pots (7 cm height × 7 cm diameter) and maintained under the same growth conditions throughout the experimental period.

### Flowering time analysis

Flowering time was determined by counting the total number of rosette leaves at bolting, excluding cotyledons ^11^. The Col-0, mutant, and transgenic plants were grown in parallel in the same tray to minimize variation in growth conditions. For flowering assays on 1/2 Murashige and Skoog (MS) media, surface-sterilized seeds were sown and grown vertically at 22 ± 1°C under a 16 h light/8 h dark cycle for 21 days.

### RNA extraction and qRT-PCR analysis

Total RNA was extracted from 200 mg of liquid nitrogen-ground samples using a series of reagents, including Trizol, chloroform, isopropanol, and ethanol. qRT-PCR was performed according to the protocol reported by Das et al. ^58^. Gene expression levels were normalized using the 2^−ΔΔCt^ method, with *P* < 0.05 considered significant. *UBQ5* served as the reference gene. Primers used in the current study are listed in Supplementary Table 1.

### Visualization of the expression levels using the Arabidopsis eFP Browser

Expression patterns of cytokinin responses were retrieved and visualized from the *Arabidopsis* eFP Browser ^59^.

### Cytokinin treatments and measurement of root length

For post-germination root growth assays, five-day-old seedlings were initially grown on vertically positioned plates containing 1/2 MS medium and then transferred to fresh growth medium supplemented with or without trans-Zeatin (*t*Z). The primary root growth was measured from photographs captured 7 days after transfer and analyzed using ImageJ software.

### Measurement of S-nitrosothiols

S-nitrosothiols (SNO) concentrations were quantified using a Sievers NOA-280i Nitric Oxide Analyser (Estero, FL, USA), following the method described by Hussain et al. ^60^. Briefly, fresh samples (200 mg) of fresh leaves were powdered in liquid nitrogen and then homogenized with 1 mL of 1× PBS buffer (pH 7.4). The homogenized solutions were centrifuged at 14,000 rpm and 4°C for 10 min to collect the supernatant. The protein content of each sample was measured using the Bradford assay. 300 µL of Coomassie dye reagent was added to 10 μL of fresh supernatant in a 96-well plate, and the mixture was mixed thoroughly by pipetting. The absorbance of the sample was measured at 595 nm using a UV spectrophotometer (Thermo Scientific Multiskan GO Microplate Spectrophotometer, Finland). Subsequently, the freshly extracted supernatant (100 μL) was introduced into the reaction vessel of the analyzer containing a CuCl/cysteine reducing agent, and the peak values were recorded. A CysNO-mediated standard curve was used to determine the SNO content, which was expressed as nanomoles per microgram of protein.

### Promoter prediction

The potential promoter binding sites of the RAP2.6 TF were predicted and analyzed using PlantPAN 4.0 ^61^, following the previous methodology ^62^.

### Statistical analysis and reproducibility

All statistical analyses, including the Student’s two-tailed *t*-test and analysis of variance (ANOVA) with Tukey’s test, were performed using “ggplot2”, “ggpubr”, “dplyr”, “agricolae”, and “tibble” in RStudio (R 4.4.3). For the purpose of reproducibility, three independent biological replicates were utilized in qRT-PCR. The sample size and the number of experimental repetitions were detailed in the figure legend.

## Acknowledgements

We are grateful to the Global Korea Scholarship (GKS) of the Government of the Republic of Korea for supporting Ashim Kumar Das. We acknowledge the help from previous lab members, Dr. Qari Muhammad Imran and Dr. Noreen Falak, in generating the double mutant.

## Author contributions

A.K.D. conceptualized the study, performed the experimental work, analyzed the data, and wrote the manuscript; M.G.M. supervised and edited the manuscript; D.-S.L. helped in data curation; B.-W.Y. supervised the study and contributed to the resources and funding.

## Data availability

All data supporting the findings of this study are available from Dr. Byung-Wook Yun upon reasonable request.

## References

1 Srikanth, A. & Schmid, M. Regulation of flowering time: all roads lead to Rome. Cell. Mol. Life Sci. 68, 2013–2037 (2011). 10.1007/s00018-011-0673-y

2 Cho, L.-H., Yoon, J. & An, G. The control of flowering time by environmental factors. The Plant Journal 90, 708–719 (2017). 10.1111/tpj.13461

3 Levy, Y. Y. & Dean, C. The Transition to Flowering. The Plant Cell 10, 1973–1989 (1998). 10.1105/tpc.10.12.1973

4 Kazan, K. & Lyons, R. The link between flowering time and stress tolerance. J. Exp. Bot. 67, 47–60 (2016). 10.1093/jxb/erv441

5. Golembeski, G. S. & Imaizumi, T. Photoperiodic Regulation of Florigen Function in Arabidopsis thaliana. The Arabidopsis Book (2015).

6 Goralogia, G. S. et al. CYCLING DOF FACTOR 1 represses transcription through the TOPLESS co-repressor to control photoperiodic flowering in Arabidopsis. The Plant Journal 92, 244–262 (2017). 10.1111/tpj.13649

7 Campos-Rivero, G. et al. Plant hormone signaling in flowering: An epigenetic point of view. J. Plant Physiol. 214, 16–27 (2017). 10.1016/j.jplph.2017.03.018

8 Izawa, T. What is going on with the hormonal control of flowering in plants? The Plant Journal 105, 431–445 (2021). 10.1111/tpj.15036

9 Schippers, J. H. M., Foyer, C. H. & van Dongen, J. T. Redox regulation in shoot growth, SAM maintenance and flowering. Curr. Opin. Plant Biol. 29, 121–128 (2016). 10.1016/j.pbi.2015.11.009

10 He, Y. et al. Nitric Oxide Represses the Arabidopsis Floral Transition. Science 305, 1968–1971 (2004). doi:10.1126/science.1098837

11 Kwon, E. et al. AtGSNOR1 function is required for multiple developmental programs in Arabidopsis. Planta 236, 887–900 (2012). 10.1007/s00425-012-1697-8

12 Zhu, W. et al. Nitric oxide delays floral transition in Arabidopsis by inhibiting histone deacetylases HDA5 and HDA6. The Plant Journal 123, e70379 (2025). 10.1111/tpj.70379

13 Yamasaki, H. & Sakihama, Y. Simultaneous production of nitric oxide and peroxynitrite by plant nitrate reductase: in vitro evidence for the NR-dependent formation of active nitrogen species. FEBS Lett. 468, 89–92 (2000). 10.1016/S0014-5793(00)01203-5

14 Rockel, P., Strube, F., Rockel, A., Wildt, J. & Kaiser, W. M. Regulation of nitric oxide (NO) production by plant nitrate reductase in vivo and in vitro. J. Exp. Bot. 53, 103–110 (2002). 10.1093/jexbot/53.366.103

15 Guo, F.-Q., Okamoto, M. & Crawford, N. M. Identification of a Plant Nitric Oxide Synthase Gene Involved in Hormonal Signaling. Science 302, 100–103 (2003). doi:10.1126/science.1086770

16 Moreau, M., Lee, G. I., Wang, Y., Crane, B. R. & Klessig, D. F. AtNOS/AtNOA1 Is a Functional Arabidopsis thaliana cGTPase and Not a Nitric-oxide Synthase*. J. Biol. Chem. 283, 32957–32967 (2008). 10.1074/jbc.M804838200

17 Lindermayr, C. & Durner, J. S-Nitrosylation in plants: Pattern and function. Journal of Proteomics 73, 1–9 (2009). 10.1016/j.jprot.2009.07.002

18 Feng, J., Chen, L. & Zuo, J. Protein S-Nitrosylation in plants: Current progresses and challenges. Journal of Integrative Plant Biology 61, 1206–1223 (2019). 10.1111/jipb.12780

19 Das, A. K. et al. The Central Role of GSNOR: Decoding Nitric Oxide Signaling for Crop Stress Tolerance. International Journal of Molecular Sciences 26, 11486 (2025). 10.3390/ijms262311486

20 Huang, Y. et al. An ethylene-responsive transcription factor and a flowering locus KH domain homologue jointly modulate photoperiodic flowering in chrysanthemum. Plant, Cell Environ. 45, 1442–1456 (2022). 10.1111/pce.14261

21 Shim, Y. et al. The AP2/ERF transcription factor LATE FLOWERING SEMI-DWARF suppresses long-day-dependent repression of flowering. Plant, Cell Environ. 45, 2446–2459 (2022). 10.1111/pce.14365

22 Nakano, T., Suzuki, K., Fujimura, T. & Shinshi, H. Genome-Wide Analysis of the ERF Gene Family in Arabidopsis and Rice. Plant Physiol. 140, 411–432 (2006). 10.1104/pp.105.073783

23 Wang, Z. et al. Identification and characterization of COI1-dependent transcription factor genes involved in JA-mediated response to wounding in Arabidopsis plants. Plant Cell Rep. 27, 125–135 (2008). 10.1007/s00299-007-0410-z

24 Zhu, Q. et al. The Arabidopsis AP2/ERF transcription factor RAP2.6 participates in ABA, salt and osmotic stress responses. Gene 457, 1–12 (2010). 10.1016/j.gene.2010.02.011

25 Krishnaswamy, S., Verma, S., Rahman, M. H. & Kav, N. N. V. Functional characterization of four APETALA2-family genes (RAP2.6, RAP2.6L, DREB19 and DREB26) in Arabidopsis. Plant Mol. Biol. 75, 107-127 (2011). 10.1007/s11103-010-9711-7

26 Fowler, S. & Thomashow, M. F. Arabidopsis Transcriptome Profiling Indicates That Multiple Regulatory Pathways Are Activated during Cold Acclimation in Addition to the CBF Cold Response Pathway. The Plant Cell 14, 1675–1690 (2002). 10.1105/tpc.003483

27 Ali, M. A., Abbas, A., Kreil, D. P. & Bohlmann, H. Overexpression of the transcription factor RAP2.6 leads to enhanced callose deposition in syncytia and enhanced resistance against the beet cyst nematode Heterodera schachtiiin Arabidopsis roots. BMC Plant Biol. 13, 47 (2013). 10.1186/1471-2229-13-47

28 Zhu, Y. et al. CDK8 is associated with RAP2.6 and SnRK2.6 and positively modulates abscisic acid signaling and drought response in Arabidopsis. New Phytol. 228, 1573–1590 (2020). 10.1111/nph.16787

29 Liu, J. et al. Identification and expression analysis of ERF transcription factor genes in petunia during flower senescence and in response to hormone treatments. J. Exp. Bot. 62, 825–840 (2011). 10.1093/jxb/erq324

30 Yin, X. et al. PfERF106, a novel key transcription factor regulating the biosynthesis of floral terpenoids in Primula forbesii Franch. BMC Plant Biol. 24, 851 (2024). 10.1186/s12870-024-05567-7

31 He, Y. et al. Research Advances in AP2/ERF Transcription Factors in Rice Growth and Development. Plants 14, 2673 (2025). 10.3390/plants14172673

32 Li, H. et al. The AP2/ERF transcription factor TOE4b regulates photoperiodic flowering and grain yield per plant in soybean. Plant Biotechnol. J. 21, 1682–1694 (2023). 10.1111/pbi.14069

33 Gibbs, Daniel J. et al. Nitric Oxide Sensing in Plants Is Mediated by Proteolytic Control of Group VII ERF Transcription Factors. Mol. Cell 53, 369–379 (2014). 10.1016/j.molcel.2013.12.020

34 Gil, M. I., Ubeda Tomas, S., Le, H. T. & Holdsworth, M. J. Recent advances in the mechanism, transduction and function of oxygen sensing in plants. J. Exp. Bot. (2025). 10.1093/jxb/eraf515

35 Falak, N., Imran, Q. M., Hussain, A. & Yun, B.-W. Transcription Factors as the “Blitzkrieg” of Plant Defense: A Pragmatic View of Nitric Oxide’s Role in Gene Regulation. International Journal of Molecular Sciences 22, 522 (2021). 10.3390/ijms22020522

36 Hussain, A. et al. Nitric Oxide Mediated Transcriptome Profiling Reveals Activation of Multiple Regulatory Pathways in Arabidopsis thaliana. Frontiers in Plant Science Volume 7–2016 (2016). 10.3389/fpls.2016.00975

37 D’Aloia, M. et al. Cytokinin promotes flowering of Arabidopsis via transcriptional activation of the FT paralogue TSF. The Plant Journal 65, 972–979 (2011). 10.1111/j.1365-313X.2011.04482.x

38 Bartrina, I. et al. Root-derived cytokinin regulates Arabidopsis flowering time through components of the age pathway. Plant Physiol. 198, kiaf204 (2025). 10.1093/plphys/kiaf204

39 Takei, K., Yamaya, T. & Sakakibara, H. Arabidopsis CYP735A1 and CYP735A2 encode cytokinin hydroxylases that catalyze the biosynthesis of trans-zeatin. J. Biol. Chem. 279, 41866–41872 (2004). 10.1074/jbc.M406337200

40 Jeon, J., Cho, C., Lee, M. R., Van Binh, N. & Kim, J. CYTOKININ RESPONSE FACTOR2 (CRF2) and CRF3 Regulate Lateral Root Development in Response to Cold Stress in Arabidopsis. The Plant Cell 28, 1828–1843 (2016). 10.1105/tpc.15.00909

41 Streatfield, S. J. et al. The Phosphoenolpyruvate/Phosphate Translocator Is Required for Phenolic Metabolism, Palisade Cell Development, and Plastid-Dependent Nuclear Gene Expression. The Plant Cell 11, 1609–1621 (1999). 10.1105/tpc.11.9.1609

42 Shi, Y.-F. et al. Loss of GSNOR1 Function Leads to Compromised Auxin Signaling and Polar Auxin Transport. Molecular Plant 8, 1350–1365 (2015). 10.1016/j.molp.2015.04.008

43 Seligman, K., Saviani, E. E., Oliveira, H. C., Pinto-Maglio, C. A. F. & Salgado, I. Floral Transition and Nitric Oxide Emission During Flower Development in Arabidopsis thaliana is Affected in Nitrate Reductase-Deficient Plants. Plant and Cell Physiology 49, 1112–1121 (2008). 10.1093/pcp/pcn089

44 Yu, X., Sukumaran, S. & Márton, L. s. Differential Expression of the Arabidopsis Nia1 andNia2 Genes: Cytokinin-Induced Nitrate Reductase Activity Is Correlated With Increased Nia1 Transcription and mRNA Levels. Plant Physiol. 116, 1091–1096 (1998). 10.1104/pp.116.3.1091

45 Olas, J. J. & Wahl, V. Tissue-specific NIA1 and NIA2 expression in Arabidopsis thaliana. Plant Signaling & Behavior 14, 1656035 (2019). 10.1080/15592324.2019.1656035

46 Tang, X. et al. The Regulation of Nitrate Reductases in Response to Abiotic Stress in Arabidopsis. International Journal of Molecular Sciences 23, 1202 (2022). 10.3390/ijms23031202

47 Methela, N. J. et al. Nitrate Reductase Genes AtNIA1 and AtNIA2 Confer Heat Stress Resilience via ROS Homeostasis and HSP Expression in Arabidopsis. Biomolecules 16, 415 (2026). 10.3390/biom16030415

48 Zhu, L., Liu, D., Li, Y. & Li, N. Functional Phosphoproteomic Analysis Reveals That a Serine-62-Phosphorylated Isoform of Ethylene Response Factor110 Is Involved in Arabidopsis Bolting Plant Physiol. 161, 904–917 (2013). 10.1104/pp.112.204487

49 Chen, Y., Zhang, L., Zhang, H., Chen, L. & Yu, D. ERF1 delays flowering through direct inhibition of FLOWERING LOCUS T expression in Arabidopsis. Journal of Integrative Plant Biology 63, 1712–1723 (2021). 10.1111/jipb.13144

50 Kardailsky, I. et al. Activation Tagging of the Floral Inducer FT. Science 286, 1962–1965 (1999). doi:10.1126/science.286.5446.1962

51 Yamaguchi, A., Kobayashi, Y., Goto, K., Abe, M. & Araki, T. TWIN SISTER OF FT (TSF) Acts as a Floral Pathway Integrator Redundantly with FT. Plant and Cell Physiology 46, 1175–1189 (2005). 10.1093/pcp/pci151

52 Samach, A. et al. Distinct Roles of CONSTANS Target Genes in Reproductive Development of Arabidopsis. Science 288, 1613–1616 (2000). doi:10.1126/science.288.5471.1613

53 Wigge, P. A. et al. Integration of Spatial and Temporal Information During Floral Induction in Arabidopsis. Science 309, 1056–1059 (2005). doi:10.1126/science.1114358

54 Endo, M. et al. Re-Evaluation of Florigen Transport Kinetics with Separation of Functions by Mutations That Uncouple Flowering Initiation and Long-Distance Transport. Plant and Cell Physiology 59, 1621–1629 (2018). 10.1093/pcp/pcy063

55 Mizoguchi, T. et al. Distinct Roles of GIGANTEA in Promoting Flowering and Regulating Circadian Rhythms in Arabidopsis. The Plant Cell 17, 2255–2270 (2005). 10.1105/tpc.105.033464

56 Feechan, A. et al. A central role for S-nitrosothiols in plant disease resistance. Proceedings of the National Academy of Sciences 102, 8054–8059 (2005).

57 Wilkinson, J. Q. & Crawford, N. M. Identification and characterization of a chlorate-resistant mutant of Arabidopsis thaliana with mutations in both nitrate reductase structural genes NIA1 and NIA2. Molecular and General Genetics MGG 239, 289–297 (1993). 10.1007/BF00281630

58 Das, A. K. et al. S-nitrosoglutathione enhances rice tolerance to salt-submergence by coordinating ethylene, GA, and ABA accumulation and improving ion transport. Plant Physiol. Biochem. 231, 111036 (2026). 10.1016/j.plaphy.2026.111036

59 Winter, D. et al. An “Electronic Fluorescent Pictograph” Browser for Exploring and Analyzing Large-Scale Biological Data Sets. PLOS ONE 2, e718 (2007). 10.1371/journal.pone.0000718

60. Hussain, A., Yun, B.-W. & Loake, G. J. in Nitric Oxide: Methods and Protocols (eds Alexander Mengel & Christian Lindermayr) 223-230 (Springer New York, 2018).

61 Chow, C.-N. et al. PlantPAN 4.0: updated database for identifying conserved non-coding sequences and exploring dynamic transcriptional regulation in plant promoters. Nucleic Acids Res. 52, D1569–D1578 (2024). 10.1093/nar/gkad945

62 Ilyas, A., et al. Enterobacter sp. SA187 boosts high-affinity nitrate transporters expression, ethylene signaling, and plant growth under low nitrate. New Phytol. 249, 3021–3038 (2026). 10.1111/nph.70885

