## Supplementary File for "AP2/ERF transcription factor RAP2.6 regulates early flowering in *Arabidopsis thaliana* by altering *S*-nitrosothiol levels and cytokinin responses"

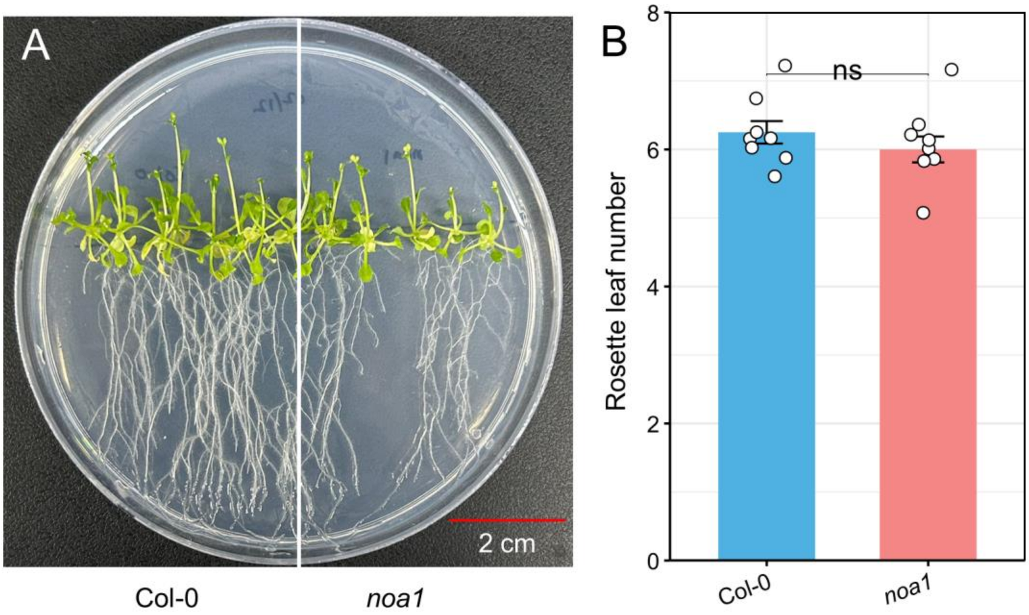


**Supplementary Fig. 1.** (A) Representative images of flowering phenotypes in Col-0 and *noa1* grown on 1/2 MS media for 21 days. (B) Number of rosette leaves at flowering in Col-0 and *noa1* mutants. The data are means ± SE ($n$ = 8 biological replicates). Asterisks indicate significant differences by Student's two-tailed $t$-test ($*P < 0.05$, $**P < 0.01$, $***P < 0.001$, $****P < 0.0001$).

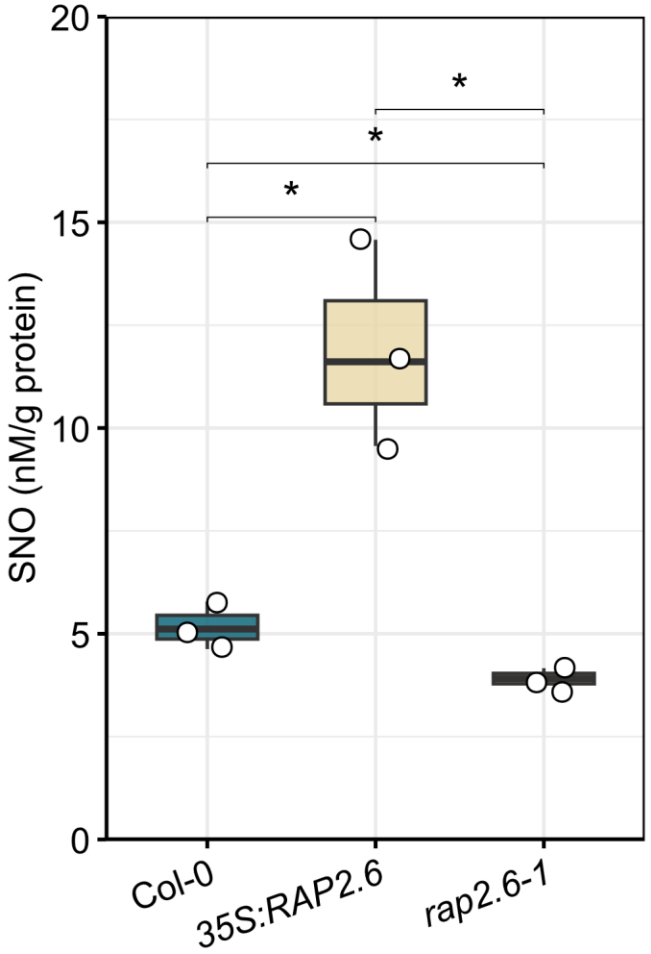


**Supplementary Fig. 2.** S-nitrosothiol (SNO) levels in Col-0, *35S::RAP2.6*, and *rap2.6-1* grown in soil. The data are means ± SE (n = 3). Asterisks indicate significant differences by Student's two-tailed *t*-test (∗$P$ < 0.05, ∗∗$P$ < 0.01, ∗∗∗$P$ < 0.001, ∗∗∗∗$P$ < 0.0001).

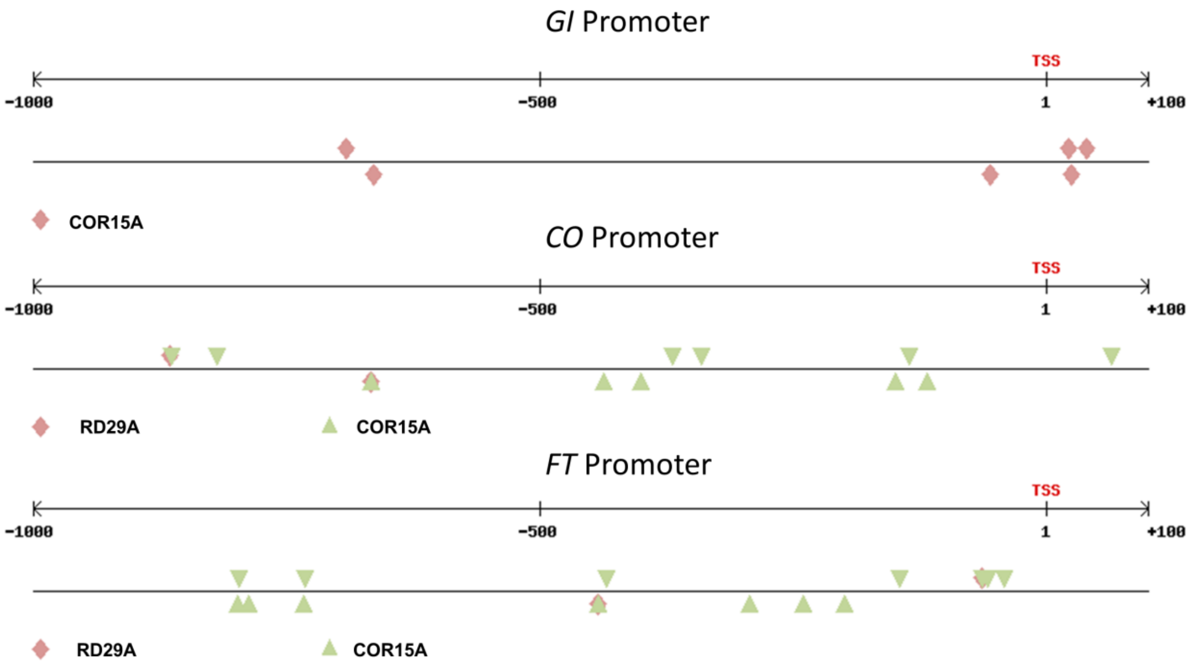


**Supplementary Fig. 3**. *In silico* analysis of the GI (AT1G22770), CO (AT5G15840), and FT (AT1G65480) promoters for potential RAP2.6 transcription factor binding sites in *cis* elements of COR15A and RD29A (Zhu et al., 2020). Promoter regions (1000 bp upstream of the transcription start site) were analyzed using PLANTPAN 4.0 under default parameters (Chow et al., 2024).

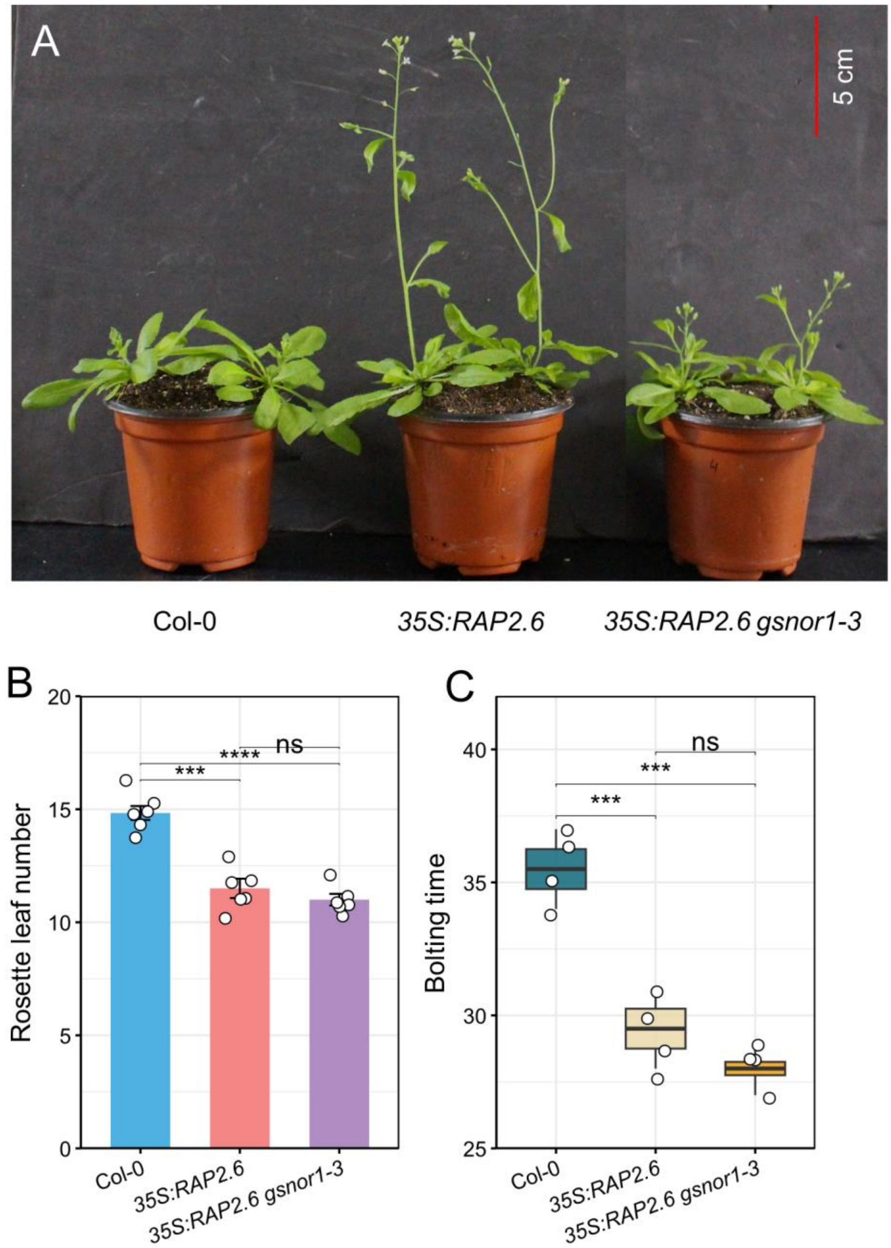


**Supplementary Fig. 4.** (A-C) Flowering phenotype, rosette leaf number, and bolting time were assessed in 38-day-old soil-grown plants of Col-0, *35S::RAP2.6*, and *35S::RAP2.6 gsnor1-3* under long-day growth conditions. The data are means ± SE ($n$ = 6 for rosette leaf number and $n$ = 4 for bolting time). Asterisks indicate significant differences by Student's two-tailed $t$-test ($*P < 0.05$, $**P < 0.01$, $***P < 0.001$, $****P < 0.0001$).

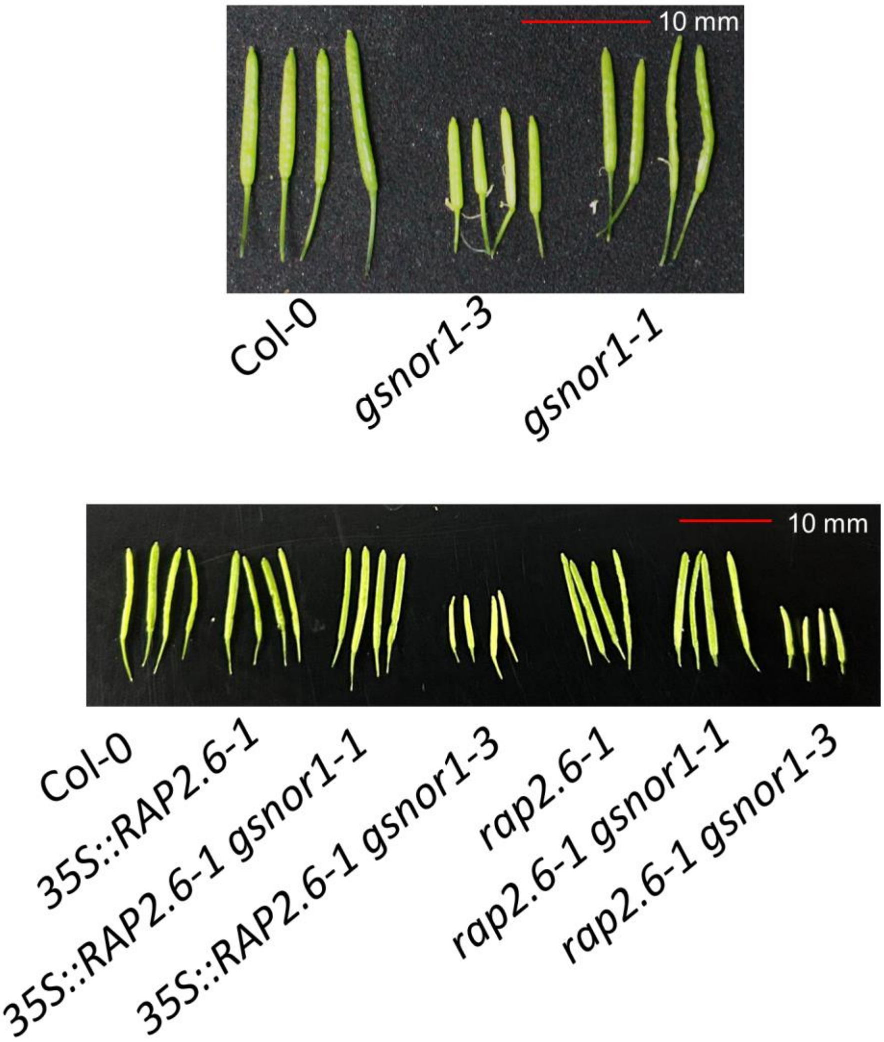


**Supplementary Fig. 5.** Representative silique phenotypes of Col-0, *gsnor1-3*, *gsnor1-1*, *35S::RAP2.6*, *35S::RAP2.6 gsnor1-1*, *35S::RAP2.6 gsnor1-3*, *rap2.6-1*, *rap2.6-1 gsnor1-1* and *rap2.6-1 gsnor1-3*.

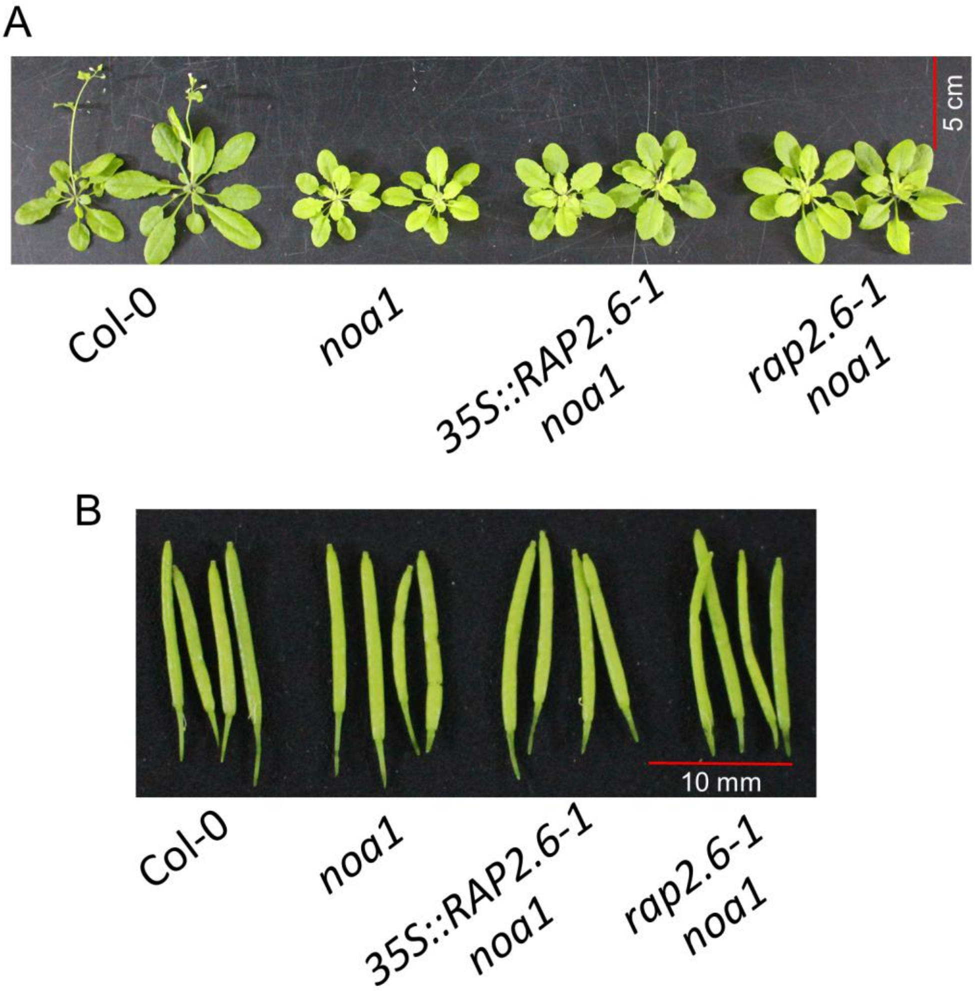


**Supplementary Fig. 6.** (A) Flowering phenotype in 38-day-old soil-grown plants of Col-0, *noa1*, *35S::RAP2.6 noa1*, and *rap2.6-1 noa1* grown in soil under long-day growth conditions. (B) Representative silique phenotypes of Col-0, *noa1*, *35S::RAP2.6 noa1*, and *rap2.6-1 noa1*.

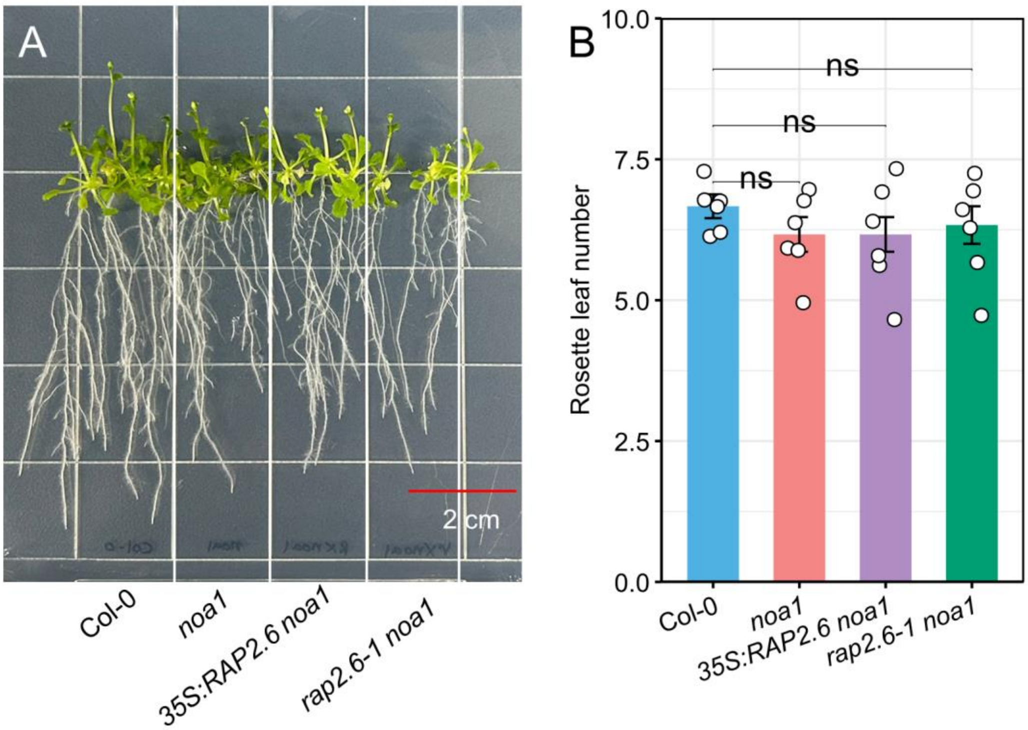


**Supplementary Fig. 7.** (A) Representative images of flowering phenotypes in Col-0, *noa1*, *35S::RAP2.6 noa1*, and *rap2.6-1 noa1* mutants grown on 1/2 MS media for 21 days. (B) Number of rosette leaves at flowering in Col-0, *noa1*, *35S::RAP2.6 noa1*, and *rap2.6-1 noa1* mutants. The data are means ± SE ($n = 6$). Asterisks indicate significant differences by Student's two-tailed *t*-test ($*P < 0.05$, $**P < 0.01$, $***P < 0.001$, $****P < 0.0001$).

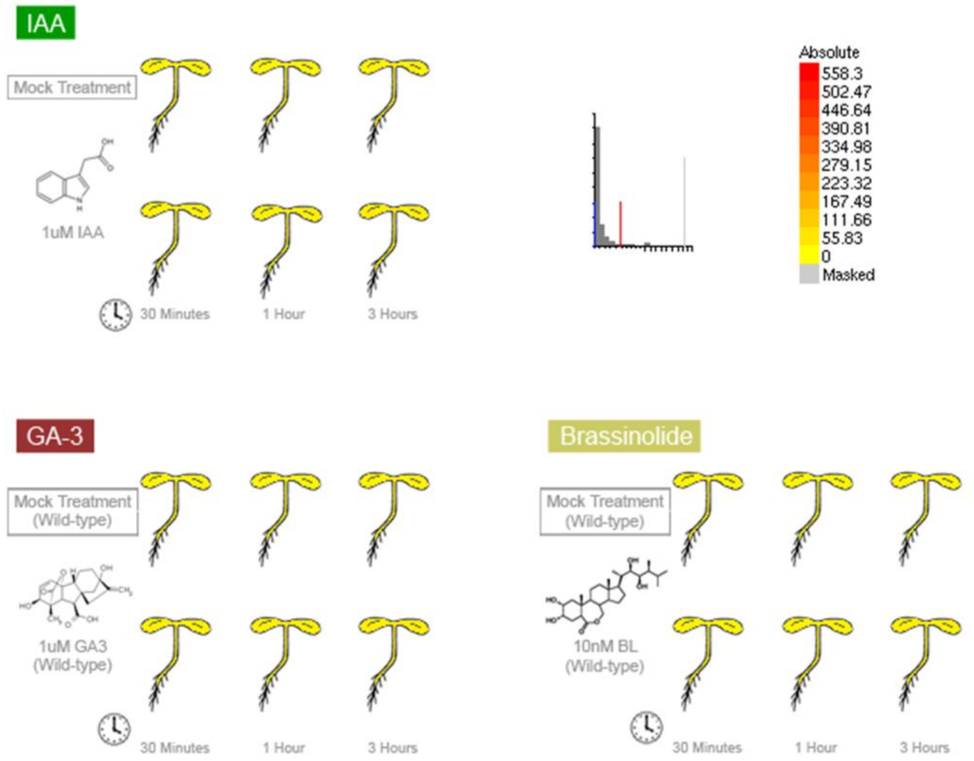


**Supplementary Fig. 8.** Visualization of the expression levels of *AtRAP2.6* in IAA, GA-3, and Brassinolide, retrieved from Arabidopsis eFP Browser (Winter et al., 2007).

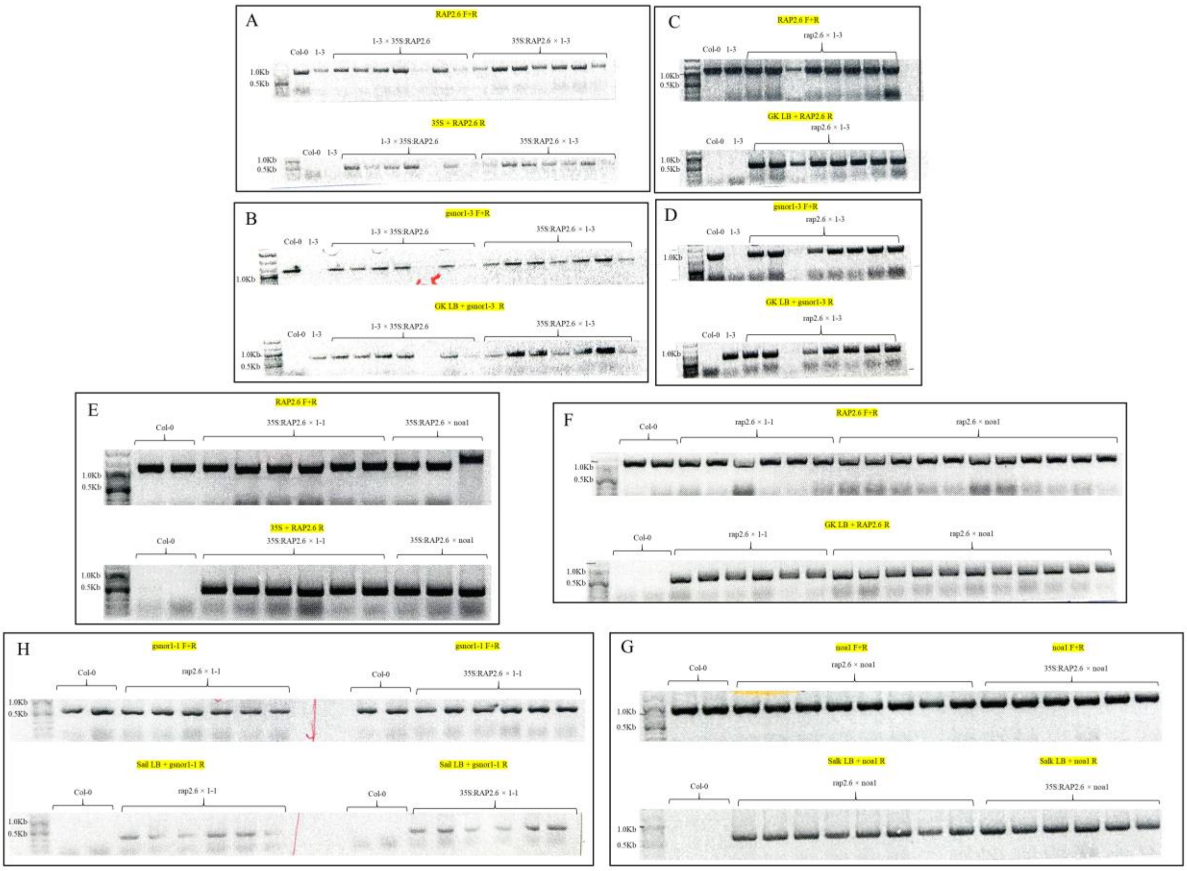


**Supplementary Fig. 9.** (A–G) Genotyping of *RAP2.6* overexpressing and knockout F1 plants after crossing with *gsnor1-3*/*gsnor1-1*/*noa1* using the respective primers. All F1 genetic epistasis combinations were heterozygous, confirming successful crossing.

**Supplementary Table 1.** List of primers used in this study.

| **For qRT-PCR** | Sequence (5' --> 3') |
|---|---|
| *AtUBQ5 F* | GGAATCGACGCTTCATCTCG |
| *AtUBQ5 R* | ACTCCTTCCTCAAACGCTGA |
| *AtCDF1 F* | AGGTGGGACCATGAGAAGTG |
| *AtCDF1 R* | TCATCTCCGAGGCTGAAACT |
| *AtFT F* | AAGTCCTAGCAACCCTCACC |
| *AtFT R* | TGAATTCCTGCAGTGGGACT |
| *AtFLC F* | AGCCAAGAAGACCGAACTCA |
| *AtFLC R* | GGGAGAGTCACCGGAAGATT |
| *AtSOC1 F* | TGCAACAAGCAGACAAGTGA |
| *AtSOC1 R* | GCATATTGGAGCTGGCGAAT |
| *AtCO F* | CACTACAACGACAATGGTTCC |
| *AtCO R* | GTCAGGTTGTTGCTCTACTG |
| *AtGI F* | CCGCCTCAAGGACTGAAATG |
| *AtGI R* | ACCACAATAGAACCCTGCGA |
| *AtCRF2 F* | CGCCGTCGTGTTAAGAAGTT |
| *AtCRF2 R* | CGCTGAAACAACCGGAGAAA |
| *AtCYP735A1 F* | TCAACCACCTCACTGTCCTC |
| *AtCYP735A1 R* | CTACCCATTTCAGCGCAGTC |
| *AtCYP735A2 F* | CCGGTCATGAGACAACCTCT |
| *AtCYP735A2 R* | ACACCATCTTGGCCACAAAC |
| **For genotyping** | Sequence (5' --> 3') |
| *AtRAP2.6-1_F* (Ali et al., 2013) | TGATGTCGGCAGTTACAAGTG |
| *AtRAP2.6-1_R* (Ali et al., 2013) | TTCCTCTAAAGCGAAGTGCTG |
| *Atgsnor1-1_F* | TATATAATGGTTCGACGCATATTT |
| *Atgsnor1-1_R* | GTCGGGTCGGGTCGTTAATA |
| *Atgsnor1-1_Sail LB* | TTCATAACCAATCTGGATACA |
| *Atgsnor1-3_F* | TATATAATGGTTCGACGCATA |
| *Atgsnor1-3_R* | CCACCAACACTCTCAACAATC |
| *Atgsnor1-3_GABI LB* | ATATTGACCATCATACTCATT |
| *Atnos1(Crawford primer)_F* | CCTGCAACACAACAAGTTGACG |
| *Atnos1(Crawford primer)_R* | GGGCGGATATAACCGTCTCCA |
